# A neuron model with unbalanced synaptic weights explains asymmetric effects of ketamine in auditory cortex

**DOI:** 10.1101/2022.06.12.495822

**Authors:** Luciana López-Jury, Francisco García-Rosales, Eugenia González-Palomares, Johannes Wetekam, Julio C. Hechavarria

**Affiliations:** Institute for Cell Biology and Neuroscience, Goethe University, 60438, Frankfurt am Main, Germany

## Abstract

Although new advances in neuroscience allow the study of vocal communication in awake animals, substantial progress in the processing of vocalizations has been made from brains of anaesthetized preparations. Thus, understanding how anaesthetics affect neuronal responses is of paramount importance. Here, we used electrophysiological recordings and computational modelling to study how the auditory cortex of bats responds to vocalizations under anaesthesia and in wakefulness. We found that multifunctional neurons that process echolocation and communication sounds were affected by ketamine anaesthesia in a manner that could not be predicted by known anaesthetic effects. In wakefulness, acoustic contexts (preceding echolocation or communication sequences) led to stimulus-specific suppression of lagging sounds, accentuating neuronal responses to sound transitions. However, under anaesthesia, communication contexts (but not echolocation) led to a global suppression of responses to lagging sounds. Such asymmetric effect was dependent on the frequency composition of the contexts and not on their temporal patterns. We constructed a neuron model that could replicate the data obtained *in vivo*. In the model, anaesthesia modulates spiking activity in a channel-specific manner, decreasing responses of cortical inputs tuned to high-frequency sounds and increasing adaptation in the respective cortical synapses. Combined, our findings obtained *in vivo* and *in silico* reveal that ketamine anaesthesia does not reduce uniformly the neurons’ responsiveness to low and high frequency sounds. This effect depends on combined mechanisms that unbalance cortical inputs and ultimately affect how auditory cortex neurons respond to natural sounds in anaesthetized preparations.

## Introduction

In the past decades, substantial progress has been made in the field of sensory processing of vocalizations. Work on anaesthetized animals has contributed tremendously to the current body of knowledge on how behaviourally-relevant sounds are represented in the brain (Suga *et al*., 1978; Margoliash & Fortune, 1992; Hsu *et al*., 2004; Schnupp *et al*., 2006; Liu & Schreiner, 2007; Grimsley *et al*., 2012). Yet, our understanding of how anaesthetics influence responses to sounds is limited. In this article, we combined computational modelling and *in vivo* electrophysiological recordings from awake and anaesthetized bats (species: *Carollia perspicillata*) to study the effects of ketamine-xylazine (KX) anaesthesia on the processing of conspecifics vocalizations.

The mixture of KX is widely used in electrophysiological studies in mammals (Fig. 1) to ensure the absence of pain and the animal’s unconsciousness. However, it can induce important changes in cortical brain activity. Ketamine mainly affects glutamatergic transmission inhibiting NMDA receptors (Sorrenti *et al*., 2021). Its effect in the cortex is characterized by increasing low-frequency oscillations and reducing ongoing ‘spontaneous’ activity (Vahle-Hinz & Detsch, 2002). However, the effects of KX on stimulus-evoked activity are difficult to generalize across cortical neurons, particularly to complex sounds such as vocalizations (Syka *et al*., 2005; Karino *et al*., 2016).

**Figure 1.**
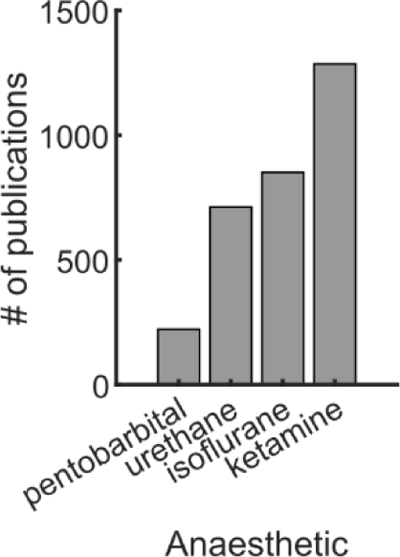
Number of publications in neuroscience using different anaesthetics in the last 10 years. Based on results from the *PubMed* search engine using the keywords combination: {‘pentobarbital’, ‘urethane’, ‘isoflurane’, ‘ketamine’} combined with the keyword ‘neuron’.

Bats represent a good mammalian model to study vocal communication (Prat *et al*., 2016; Vernes, 2017). In addition to their own sonar pulses, that allow bats to navigate in the dark, bats are constantly processing echolocation pulses and social calls from conspecifics (Fenton, 2003). These two type of vocalizations differ in terms of their frequency composition, social calls often have lower fundamental frequencies than echolocation (Pfalzer & Kusch, 2003). In the auditory cortex of bats there are specialized neurons that exhibit multiple forms of selectivity for both types of signals, echolocation and communication, suggesting a multifunction theory of cortical processing (Suga, 1994). Bat multifunctional neurons have been well characterized in terms of their location within the auditory cortex, responses to pure tones, and even to natural sounds (Ohlemiller *et al*., 1996; Esser *et al*., 1997; Kanwal, 1999; Razak *et al*., 1999; Washington & Kanwal, 2008). It has been shown that multifunctional neurons increase their discrimination between echolocation and communication calls when the vocalizations are preceded by natural acoustic contexts (Lopez-Jury *et al*., 2021). Multifunctional neurons present a good opportunity to investigate the effects of anaesthesia on sensory processing of vocalizations due to (i): they constitute a well characterized neuronal population; (ii) they are thought to play an important role for communication in complex acoustic environments; and (iii) the synaptic mechanisms underlying the effects of anaesthesia can be studied *in silico*, since a neuron model that explains responses to individual sounds and sound mixtures in the awake state is available (Lopez-Jury *et al*., 2021).

The main aim of this study was to determine if echolocation and communication acoustic contexts drive similar effects on lagging sounds in awake and KX anaesthetized bats. Previous studies have shown that ketamine reduces responses to vocalizations (Syka *et al*., 2005; Karino *et al*., 2016) and increases adaptation in cortical regions (Rennaker *et al*., 2007; Ter-Mikaelian *et al*., 2007). Considering these findings, we modified the neuron model previously constructed for awake data and simulated the expected KX effects. We called this first *in silico* experiment “naïve modelling”. Naïve modelling predicted that known effects of KX would have significant effects on the context-mediated modulation of neural responses, independent on the type of the context that precedes the target sounds (echolocation or communication). Surprisingly, electrophysiological recordings contradicted the naïve model prediction and showed that under KX, context effects were asymmetrical: only the sound responses after communication context were affected by anaesthesia. A new set of *in vivo* experiments allowed us to determine that such asymmetries were due to stimulus frequency-specific effects of anaesthesia. The naïve model was updated based on the findings obtained *in vivo*. From the updated models, the one that succeeded at reproducing all *in vivo* experimental observations was one in which anaesthesia affected only high-frequency tuned inputs and their respective synapses, creating a compensation of the effects and causing the apparent no effect of anaesthesia after echolocation context.

## Results

### Predicted effects of anaesthesia in auditory cortex (naïve modelling)

We modified a cortical neuron model of (Lopez-Jury *et al*., 2021) to investigate the effects of KX on context dependent processing of vocalizations. The published model reproduces spiking responses to acoustic transitions between echolocation and communication sounds in awake bats. It showed that in the presence of acoustic context, sequences of echolocation and communication (Fig. S1), leads to an overall response suppression of probe-evoked responses. This suppression is stimulus-specific. To study the effects of KX on these findings, we varied model parameters in a manner consistent with experimentally observed effects of KX on evoked responses to complex and natural sounds. For instance, it had been shown that KX suppresses responses to natural sounds and increases adaptation to repetitive sounds (Zurita *et al*., 1994; Syka *et al*., 2005; Rennaker *et al*., 2007; Scholes *et al*., 2015; Karino *et al*., 2016). Accordingly, we independently decreased the average firing rate of inputs in response to the context sequences and increased the magnitude of pre- and postsynaptic adaptation, relative to the parameters values fit to the awake data (Fig. 2). The mechanisms of neuronal suppression in the model are activity dependent. Therefore, a reduction in the firing rate at the input level decreased the suppression on both probe responses (context effect values closer to zero, Fig. 2A top, middle). However, the effect size was largest in the matching probes (e.g. communication probe following communication context). Consequently, the stimulus-specific suppression index (s.s.s.), defined as the difference between context effects on matching and mismatching probes, decreased (Fig. 2A bottom). On the other hand, an increment of adaptation had the opposite outcome: it increased context effect on matching probes, in the case of presynaptic adaptation (Fig. 2B top, middle) and it increased context effect on both probes, in the case of postsynaptic adaptation (Fig. 2C top, middle). While the unbalanced effect on probe responses observed increasing presynaptic adaptation led to an increment of s.s.s. (Fig. 2B bottom), the general effect of postsynaptic adaptation had no repercussions on s.s.s. (Fig. 2C bottom).

**Figure 2.**
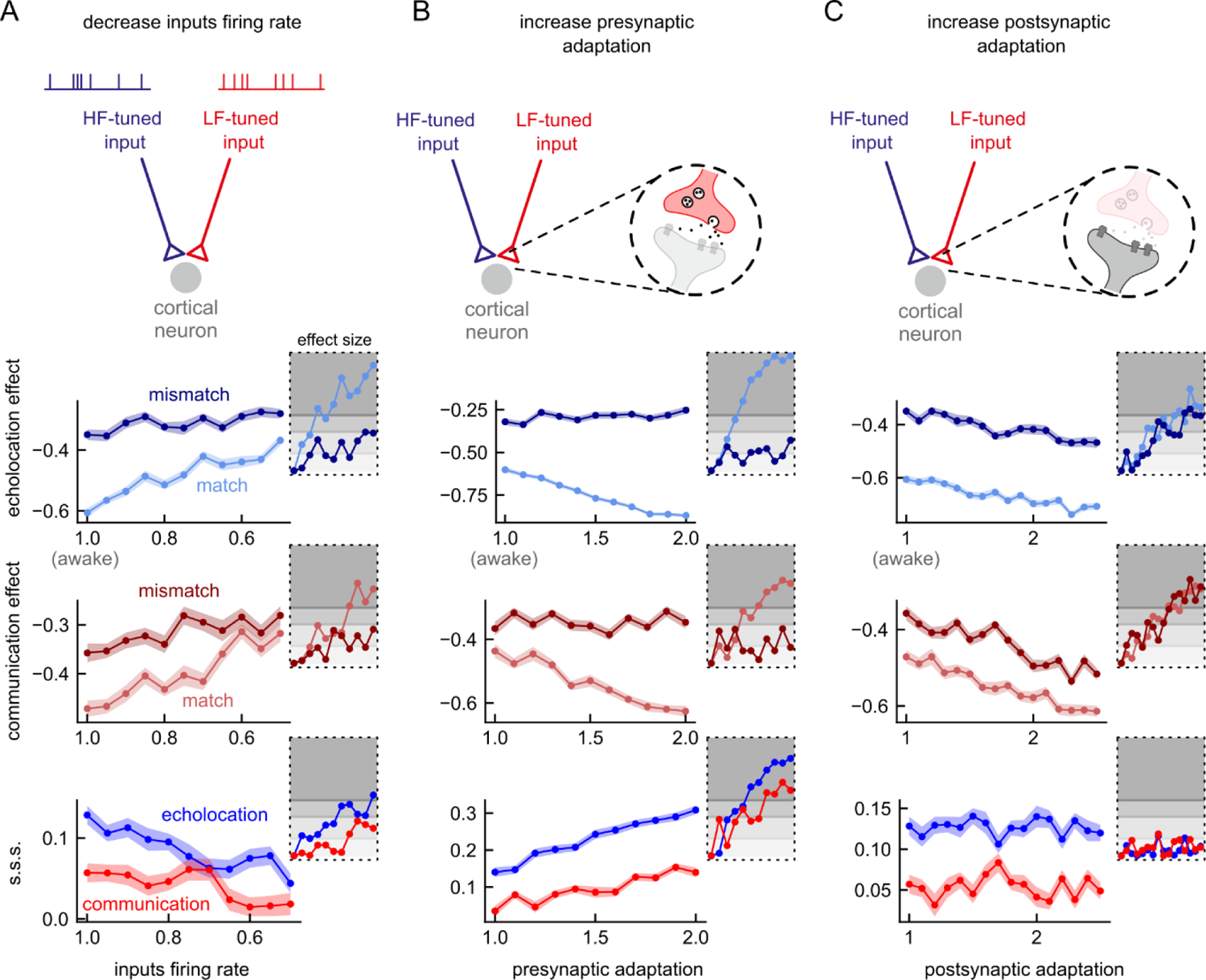
Effects of anaesthesia on a model for acoustic context effects on cortical neuron responses to natural sounds. Diagrams on top illustrate possible effects of ketamine-xylazine on a neuron model that explains acoustic context effect in the auditory cortex of awake bats (López-Jury et al. 2021). **A**. The decrease of the cortical input firing rate was implemented by systematically decreasing the average input rate parameter, ν, by a factor of 1.0 (awake model) to 0.5 in steps of 0.05. First row shows the context effect after echolocation context for each simulation (n=50) for the mismatching and the matching probes. Second row shows the context effect after communication context. Third row shows the stimulus-specific suppression index (s.s.s.) obtained from the same simulations after echolocation context (in blue) and communication context (in red). Insets indicate the effect size (Cliff’s Delta) of each simulation for each parameter compared against the respective awake model (i.e. factor = 1.0). The different degrees of grey indicate the different interpretation of effect size: negligible, small, medium and large. **B**. The increment of presynaptic adaptation was implemented by systematically increasing the parameter Δs, by a factor of 1.0 to 2.0 in steps of 0.1. **C**. The increment of postsynaptic adaptation was implemented by systematically increasing the adaptive threshold time constant, τth, by a factor of 1.0 to 2.5 in steps of 0.1. Note that presynaptic adaptation accentuated differences between probe-responses, while postsynaptic adaptation left them unchanged.

As expected, all the models of anaesthesia showed a decrement in the spiking activity in response to natural sounds compared to the awake model. In general, the size of the effects was similar between echolocation and communication or slightly larger for echolocation context than communication (insets in Fig. 2).

Next, we ran several simulations co-varying two parameters to determine how the interactions between two effects of anaesthesia influence the outcome of the models in comparison to the awake model (Fig. 3, Fig. S2). Considering that the reduction of inputs firing rate and the enhancement of adaptation have opposite consequences on the context effects in the model, we hypothesized that the combination of both could compensate each other and lead to an unchanged outcome. Indeed, the results of the simulations decreasing input activity as well as increasing presynaptic adaptation showed a region of parameters wherein changes in the context effect on matching probes and in the s.s.s. was negligible relative to the awake model (Fig. 3A, pale diagonal in first and third column). In the same simulations, the effect on the mismatching probe was generally a reduction of the context effect (positive values of Δ c.e., Fig. 3A middle column). Compensation of effects was also observed between inputs activity and postsynaptic adaptation (Fig. S2A). However, the compensatory region of parameters with negligible changes appeared for both probes, matching and mismatching. As expected, the increment of both type of adaptations, pre- and postsynaptic, generated increment on context effects; stronger suppression was observed on the matching probe (compared to the mismatching one) for all the range of parameters tested. In addition, the s.s.s. increased after both contexts due to presynaptic adaptation increment (Fig. S2B).

**Figure 3.**
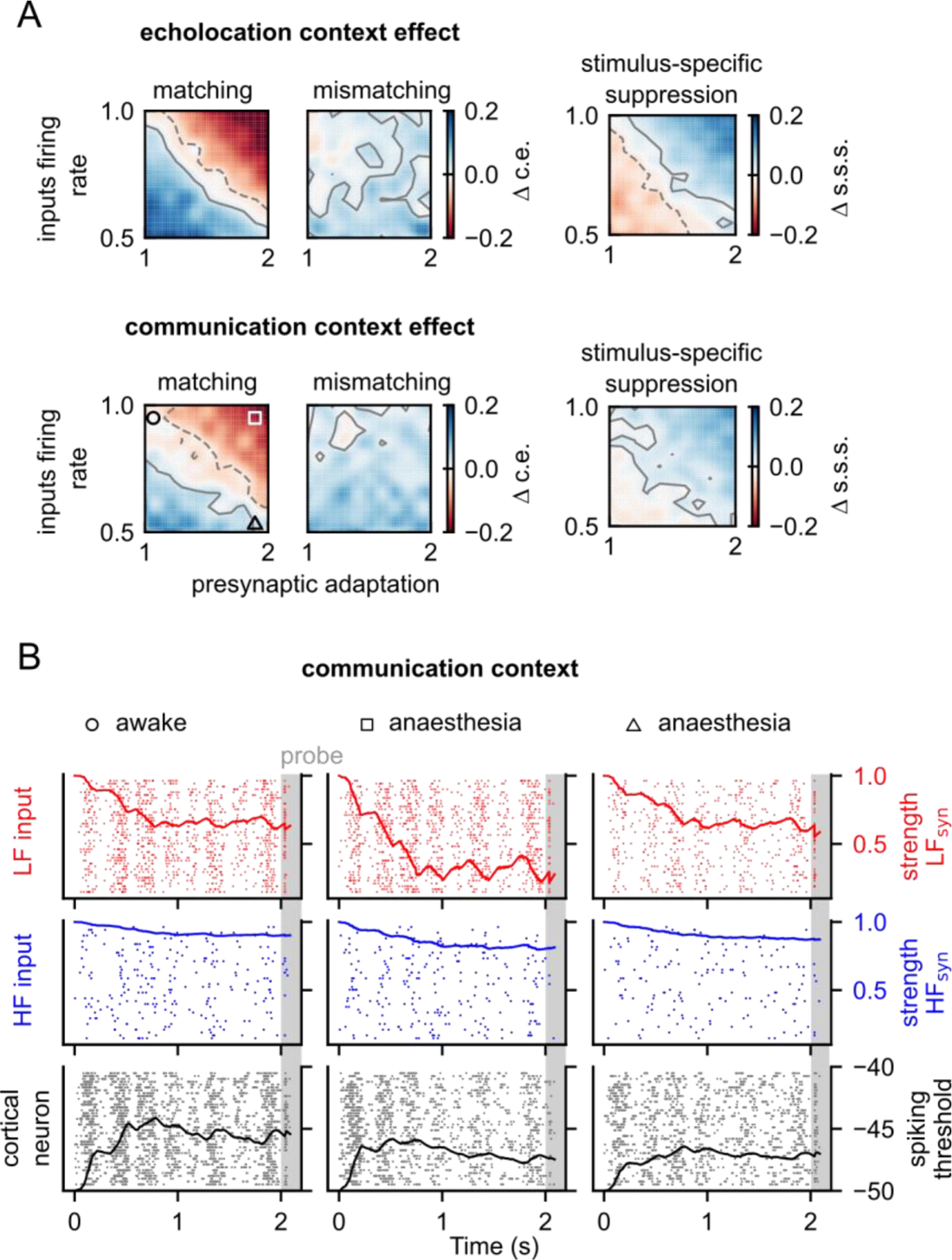
Interaction between two anaesthesia effects leads to a compensatory mechanism in the outcome of the model. **A**. Difference between context effect (left) and stimulus-specific suppression (right) when decreasing inputs firing rate and increasing presynaptic adaptation relative to the respective value obtained with the awake model. First row, after echolocation. Second row, after communication context. Contour lines in grey indicate the area with negligible effect size. Positive contours are indicated by solid lines and negative, by dashed lines. The anaesthesia effects were implemented as described in Figure 2. Note that red colours in the colormaps correspond to an increment of suppression, and blue, a decrement. **B**. Raster plots of 30 trials for the low-frequency tuned input (top), high-frequency tuned input (middle) and cortical neuron (bottom) during a simulation of communication context and matching probe for three combination of parameters: (1) no anaesthesia (circle) which corresponds to inputs firing rate factor = 1.0 and presynaptic adaptation factor = 1.0, this represents no change from the awake model; (2) under anaesthesia (square) was obtained with double of presynaptic adaptation used in the awake state and (3) under anaesthesia (triangle) was obtained with double presynaptic adaptation but half of inputs firing rate. Lines on the top rasters indicate the average strength of the synapse (*X*s) between low-frequency tuned inputs and cortical neurons along time. Lines on the middle rasters indicate the same but from the high-frequency tuned inputs. Lines on bottom rasters indicate the average spiking threshold (ωth) of the cortical neuron across trials along time.

It is important to note that both contexts, echolocation and communication, exhibited very similar patterns of change under the simulated effects of anaesthesia. To better visualize the interaction between input firing rate and adaptation, we plotted the time course of presynaptic adaptation and the respective spiking activity for each input and the cortical neurons. We compared the results for three different models during communication context simulations (Fig. 3B). In all the models, the synaptic strength associated to low-frequency inputs decreased throughout the context due to presynaptic adaptation. In the awake model, at the end of the context the synaptic strength was approximately 0.4 (Fig. 3B left column). This value depends on the total amount of spikes evoked by the context as well as on the amount of presynaptic depression per spike. In a model in which the amount of presynaptic depression per spike is higher, but the input activity is constant, the synaptic strength reached even lower values at the end of the context, approximately 0.2 (Fig. 3B middle column). Finally, decreasing the inputs spiking in the presynaptic depression increased model compensated the effect of strong spike-dependent depression by reducing the total amount of spikes in the inputs (Fig. 3B right column). The strength of the synapse at the end of the context is then similar in magnitude to that observed in the awake model (∼0.4).

To summarize this section, modifying a model based on awake data to simulate the effects of KX on neuronal activity resulted in: (i) decreased spiking in the cortical neurons independently of the type of context the animals listen to and, (ii) interactions between KX effects that could lead to “compensatory regions” in the model’s output. In such compensatory regions context effect and stimulus specific suppression appear unchanged when compared to the awake state.

### Effects of anaesthesia in in vivo preparations

To assess the effect of KX on context dependent processing of vocalizations experimentally, we performed single neuron recordings in AC of anaesthetized bats and compared the neuronal responses to the same paradigm of stimulation used in a previous study in awake animals (Lopez-Jury *et al*., 2021). Our goal was to validate one of the model predictions and thus, determine the mechanisms underlying the KX effects on evoked responses to natural sounds. We recorded a total of 107 units from high-frequency fields of the AC under anaesthesia. As expected, several neurons presented multi-peaked frequency tuning curves (n=62, Fig. S3A-B). This tuning shape has been linked to high-frequency areas in *C. perspicillata* in previous studies (Hagemann *et al*., 2010; Lopez-Jury *et al*., 2020). Here, we focus our analysis only in a subset of neurons characterized electrophysiologically as ‘equally responsive’ neurons to both sound categories: echolocation pulse and communication syllable (n = 46, Fig. S3C), following the same criteria used to describe context effects in awake bats (Lopez-Jury *et al*., 2021).

The awake and anaesthetized preparations presented similar physiological properties, such as tuning to pure tones (iso-level frequency tuning curves) and responses to natural sounds (Fig. S3D-F). Two representative examples from each preparation are given in Fig. 4. In both cases, neurons classified as “equally responsive” to both probe sounds in the absence of context, exhibited strongest responses to mismatching probe sounds after 60 ms of the end of the navigation context (Fig. 4 top). However, a different outcome was observed after the communication context. Although the response to the mismatching probe was stronger than to the matching probe in the awake example neuron, the responses under anaesthesia were strongly suppressed to both matching and mismatching probes (Fig. 4 bottom). Only the communication context under anaesthesia presented a negative value of s.s.s., indicating that the response to the matching probe was less suppressed than to the mismatching probe relative to those in isolation.

**Figure 4.**
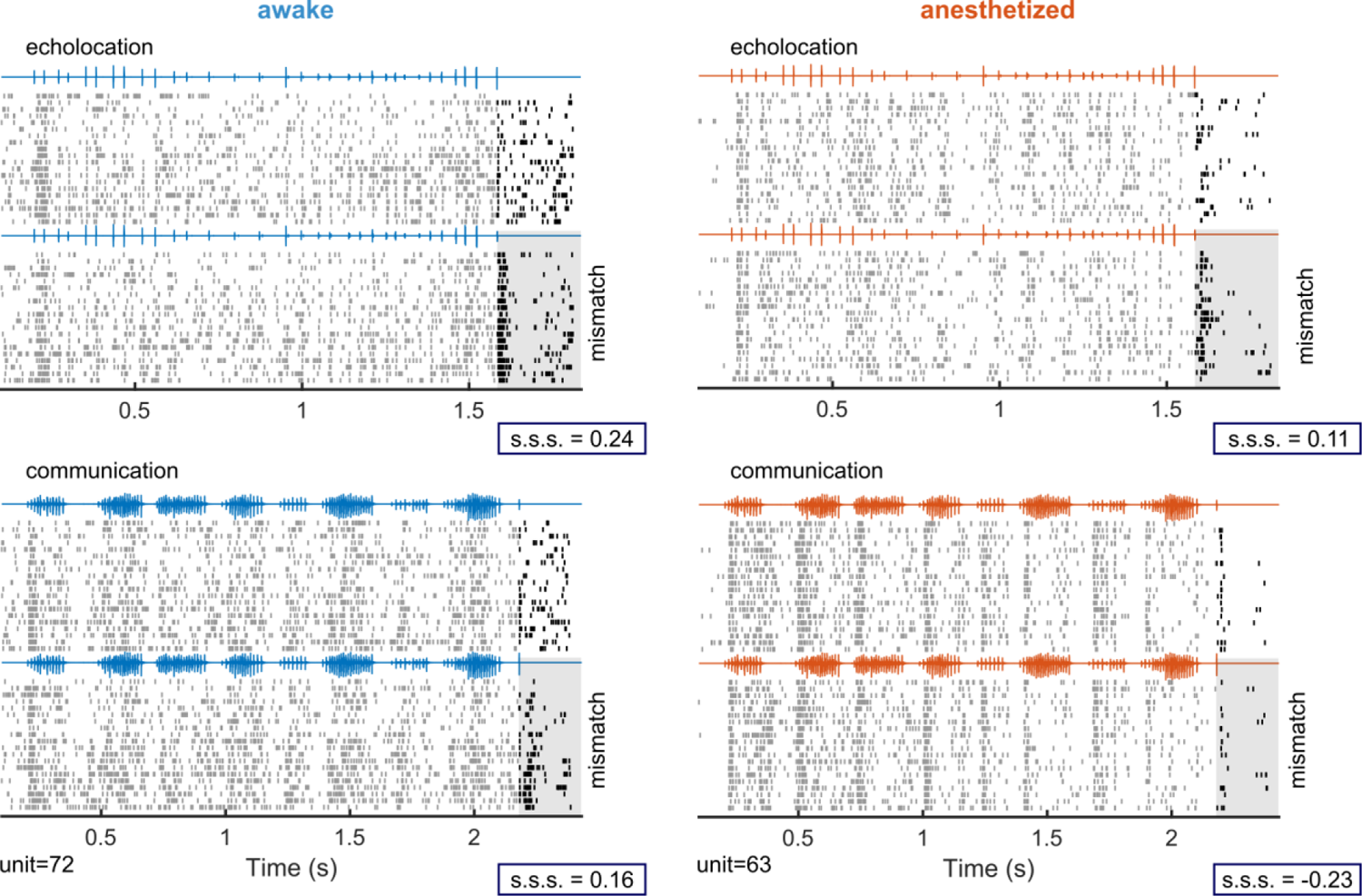
Examples of effects of anaesthesia on cortical neurons spiking for acoustic context effects on responses to natural sounds. Two example units illustrating cortical responses to natural sounds in awake (left) and anesthetized (right) bats. Top, spiking responses to a sequence of echolocation pulses (context) followed by a matching probe sound (top 20 trials) and by a mismatching probe sound (bottom 20 trials). Probe sounds correspond to a single echolocation pulse and a communication call, respectively. Bottom, the responses of the same neurons to a communication sequence followed by matching and mismatching probe sounds, i.e., a single communication call and by an echolocation pulse, respectively. The grey square is aligned to the onset of the last call that correspond to the mismatching probe sound in both cases. For each example unit, the respective value of stimulus-specific suppression (‘s.s.s.’) per context is indicated.

While the majority of neurons recorded from awake bats showed positive s.s.s. after navigation and communication contexts, neuronal responses under anaesthesia exhibited positive values only after navigation context (Fig. 5A-B). The average s.s.s. obtained after echolocation was higher than communication in both preparations (Fig. 5C, paired test Wilcoxon signed-rank, *p* = 0.046 in awake, *p* = 0.049 in anaesthetized). However, only s.s.s. after communication under anaesthesia was not different from a distribution centred on zero, the rest of the distributions were all significantly above zero (Fig. 5C, one sample Wilcoxon signed-rank test, *p* = 6.3e-4 for echolocation-awake, *p* = 1.7e-4 for communication-awake, *p* = 7.2e-4 for echolocation-anaesthetized, *p* = 0.4 for communication-anaesthetized). To visualize the spiking responses in time under both states, we plotted the instantaneous firing rate aligned to the probe onset of neurons that corresponded to the most represented category in awake and anaesthetized preparations. Although neuronal responses to echolocation and communication probe sounds in isolation were comparable in both states (Fig. S4), after echolocation context, neurons exhibited a clear preference for the mismatching sound, i.e. communication probe (Fig. 5D-E top). Regarding activity during the echolocation context, the responses under anaesthesia were lower in comparison to those in awake state. In the case of communication context in awake animals, although there was difference between the responses to both probes, the preference for the mismatching probe was less clear than after echolocation (Fig. 5D bottom). On the other hand, for units recorded under anaesthesia, there was no difference in the magnitude of the responses to the probes, both were strongly suppressed after context, independently on the probe type (Fig. 5E bottom).

**Figure 5.**
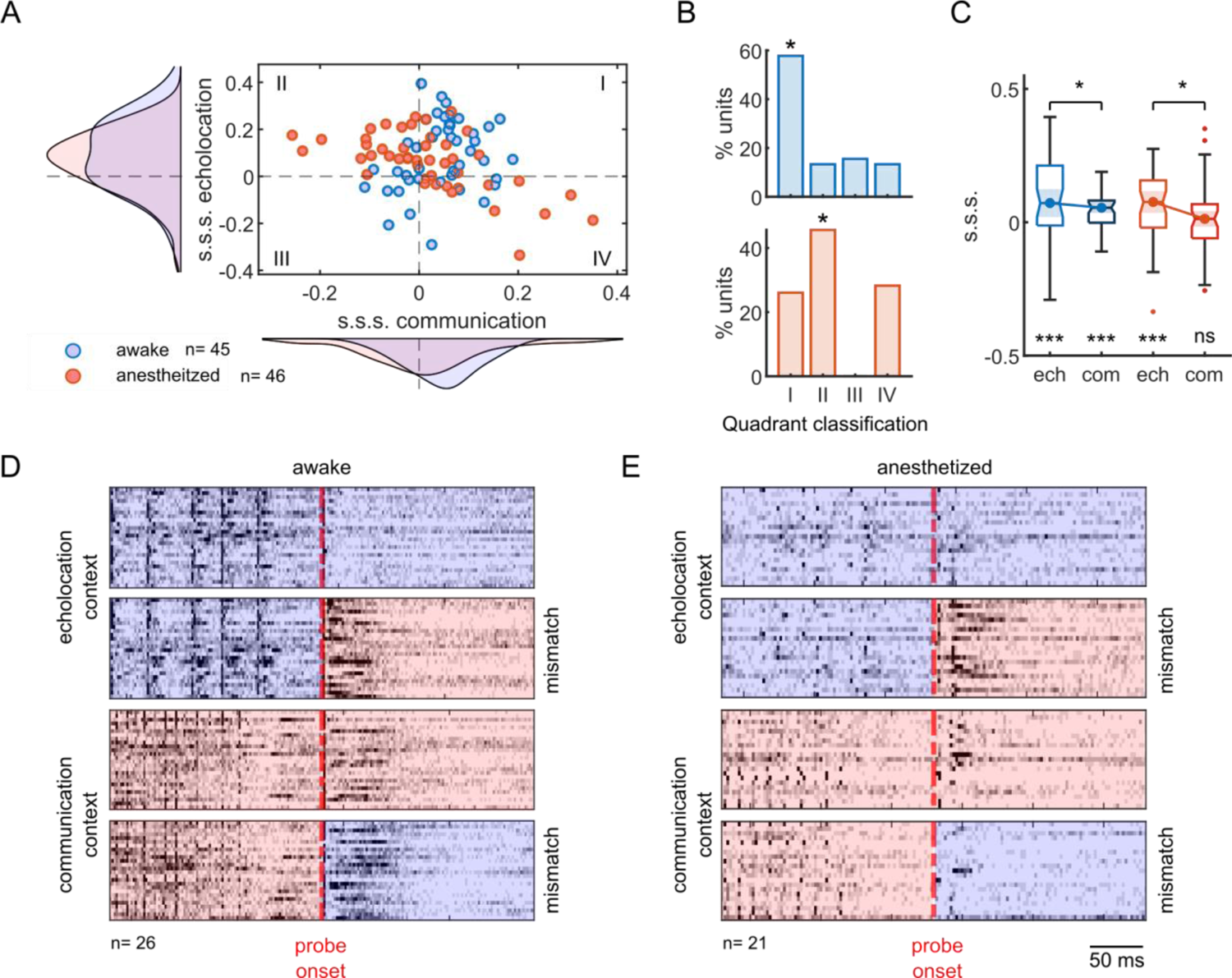
Anaesthesia decreases stimulus-specific suppression on probe-responses only after communication context. **A**. Stimulus-specific suppression indexes (s.s.s.) calculated from awake (blue) and anesthetized (red) preparations. Each circle corresponds to one unit (n=45 awake, n=46 anesthetized). The curves represent the distribution for a particular context. **B**. Percentage of units per quadrant showed in ‘a’ of units recorded in awake (top) and anesthetized animals (bottom). Asterisks show the most represented quadrant on each preparation. **C**. Boxplots of s.s.s. per context in awake and anesthetized preparations. Significance level below the boxplot indicates differences against null distribution (one sample Wilcoxon signed rank test). Above, the comparison between contexts (paired test Wilcoxon signed rank). **D**. Normalized and averaged firing rate of units that corresponded to the most representative categories in b (quadrant I for awake and quadrant II for anesthetized) aligned by probe onset. Blue background corresponds to echolocation sounds stimulation and red, to communication, either context or probe. **E**. Same than in d but for the anesthetized preparation. All the results correspond to a gap between context offset and probe onset equal to 60 ms.

In general, in both preparations, the context suppressed the responses to the probe sounds (negative values of context effect, Fig. 6A-B). In agreement with previous results quantifying s.s.s., the context effects were significantly different between probes in both states after echolocation context (Fig. 6A, Wilcoxon signed rank test, *p* = 6.3e-4 in awake, *p* = 7.2e-4 in anaesthetized) and, only in awake state, after communication (Fig. 6B, Wilcoxon signed rank test, *p* = 1.7e-4 in awake, *p* = 0.45 in anaesthetized). Using a non-paired statistical test, we compared the context effect between preparations, there were only significant differences between communication context effects. The context effect decreased for both matching and mismatching probes under anaesthesia relative to the awake treatment (Fig. 6A-B, Wilcoxon Rank-sum test, *p* = 0.33 for echolocation-matching, *p* = 0.99 for echolocation-mismatching and *p* = 0.02 for communication-matching, *p* = 0.006 for communication-mismatching).

**Figure 6.**
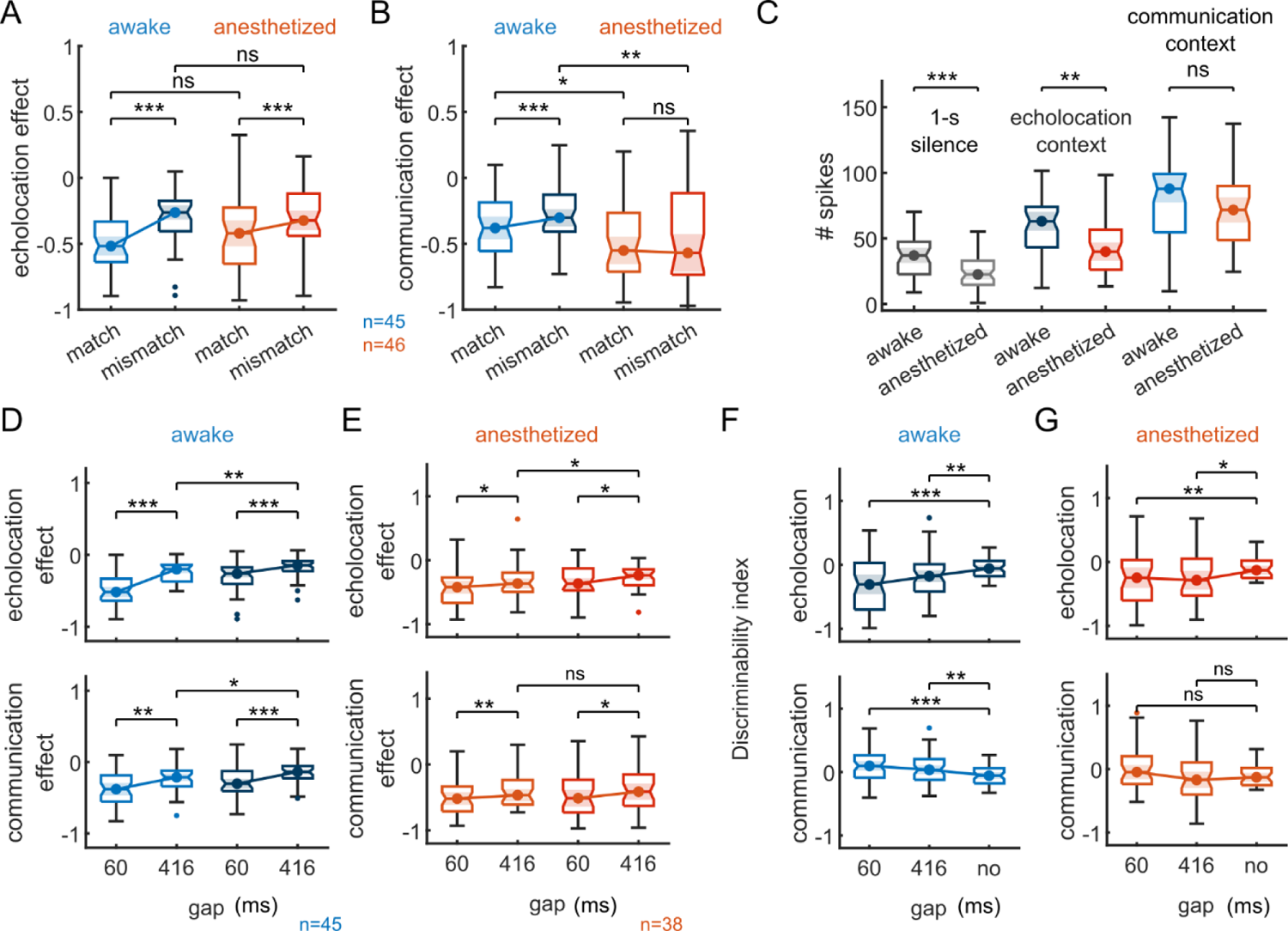
Stronger suppression on both probes after communication context explains decrement on stimulus-specific suppression. **A**. Context effect on probe-responses following an echolocation sequence. In blue, neurons recorded from awake bats (n=45); in red, from anesthetized (n=46). **B**. Context effect after communication context. **C**. Spike counts for 1-s of silence, during the echolocation context and communication context; for awake and anesthetized preparations. **D**. Top, comparisons between context effect after echolocation context with a gap of 60 ms between probe onset and context offset and a gap of 416 ms in awake animals. Bottom, same than above, but after communication context. **E**. Same than in ‘D’ but in anaesthetized animals. **F**. Discriminability index calculated after 60 and 416 ms from the offset of the echolocation context (top) and communication (bottom) and after no context in the awake preparation (n=45). **G**. Same than in ‘F’ but the neurons were recorded in anesthetized bats (n=38). Statistical tests within preparations were obtained using a paired test (Wilcoxon signed rank). Statistical differences between awake and anesthetized preparations were obtained using Wilcoxon ranksum test.

Other features such as spontaneous firing rate and evoked response by the context sequences were also compared across preparations (Fig. 6C). As expected from the effect of KX, the average spontaneous firing rate of the neurons was lower in the preparation under anaesthesia compared to awake (Fig. 6C, Wilcoxon Rank-sum test, *p* = 1.7e-4). Similarly, the evoked response during the echolocation context also decreased (Fig. 6C, Wilcoxon Rank-sum test, *p* = 0.0013). However, no significant differences were found in the evoked responses during the communication context between preparations (Fig. 6C, Wilcoxon Rank-sum test, *p* = 0.17). To corroborate the comparisons between preparations, we performed new experiments testing the effect of KX directly on the same neurons. The neurons were recorded both prior to and following anaesthesia administration. Although we recorded a reduced sample size (n=7, it was difficult to hold the same neurons for long time periods and only one neuron could be recorded in each animal) the trend already showed the same effects obtained from different preparations (Fig. S5A). The effect size between context effects calculated before and after the injection of KX was, on both probe sounds, lower after echolocation than communication context (Fig. S5A, Cliff’s delta, *d* = 0.14 for echolocation-matching, *d* = 0.18 for echolocation-mismatching, *d* = 0.30 for communication-matching and *d* = 0.32 for communication-mismatching). In addition, the same neurons showed a significant decrease in the spontaneous firing rate (Fig. S5B, Wilcoxon signed rank test, *p* = 0.01) and in the evoke response only during echolocation context (Fig. S5B, Wilcoxon signed rank test, *p* = 0.01 for echolocation and *p* = 0.15 for communication) after the KX injection. Altogether, these results replicated those obtained between different preparations.

All the results presented so far have been obtained using context-probe stimuli with a time interval of 60 ms between the end of the context and the onset of the probe. In order to explore the temporal course of the context effects, we also used a gap of 416 ms during the recordings under anaesthesia in a subset of neurons (38 out of 46). The same gap was used in the stimuli used in the neurons of the awake preparation (Lopez-Jury *et al*., 2021). In both preparations, the context effect showed significant reduction after a gap of 416 ms compared to 60 ms (Fig. 6D-E, Wilcoxon signed rank test, *p* = 6.7e-8, *p* = 2.0e-5, *p* = 0.001, *p* = 3.8e-5 for echolocation-matching, echolocation-mismatching, communication-matching and communication-mismatching respectively for awake, and *p* = 0.02, *p* = 0.03, *p* = 0.002, *p* = 0.03 for anaesthetized). Despite of the recovery of the context effects observed at longer gaps, the effect after echolocation context was still stimulus-specific, i.e. there were significant differences in context effect in matching versus mismatching probes in both states (Fig. 6D-E top, Wilcoxon signed rank test, *p* = 0.007 for awake and *p* = 0.02 for anaesthetized). On the other hand, communication context effect was stimulus specific for 416-ms gaps only in the awake preparation (Fig. 6D-E bottom, Wilcoxon signed rank test, *p* = 0.02 for awake and *p* = 0.7 for anaesthetized). This was expected considering that the effect was not stimulus-specific even 60 ms after the end of the communication sequence (see Fig. 6B). It is worth mentioning that the effect size (Cliff’s delta, *d*) of the context effects between gaps was always smaller under anaesthesia than awake (*d* = 0.46 for echolocation and *d* = 0.38 for communication in awake preparation; *d* = 0.16 for echolocation and *d* = 0.11 for communication in anaesthetized preparation), indicative of a slower recovery of the context effects under anaesthesia.

From the experiments with awake animals, the neurons increased their discriminability for the probe sounds after both contexts for both gaps compared to when there was no context (Fig. 6F, Wilcoxon signed rank test, *p* = 7.6e-5 for echolocation-60ms, *p* = 0.001 for echolocation-416ms, *p* = 1.9e-4 for communication-60ms, *p* = 0.002 for communicacion-416ms). Here, we showed the same occurred under anaesthesia, but only after echolocation context (Fig. 6G, Wilcoxon signed rank test, *p* = 0.001 for echolocation-60ms, *p* = 0.01 for echolocation-416ms, *p* = 0.2 for communication-60ms, *p* = 0.6 for communicacion-416ms).

### Asymmetric effects of anaesthesia on the processing of echolocation vs. communication sounds is linked to frequency specific effects

Anaesthesia influenced asymmetrically the cortical processing of echolocation and communication calls: KX modulated the context effects on the probe responses only after the communication context. However, the neuronal responses during the context were only affected for echolocation (summary in Fig. 7 top). None of the models presented at the beginning of this article (Figs. 2-3) predicted such asymmetrical modulation. Our results indicate that the known effects of KX implemented in our anaesthesia models cannot explain the experimental data obtained *in vivo*. We conducted several *in vivo* and modelling experiments to disentangle the possible causes of these results.

**Figure 7.**
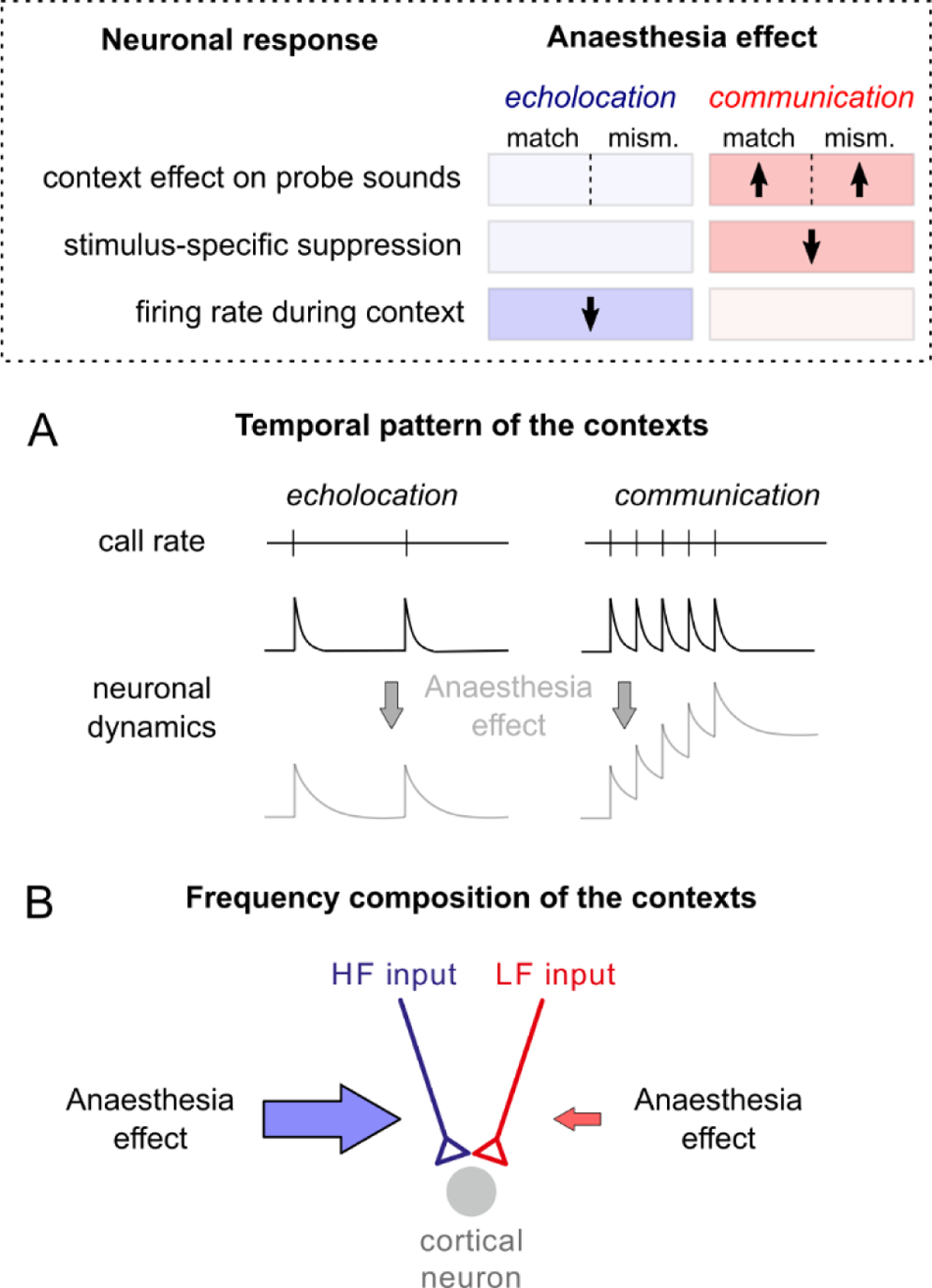
Anaesthesia modulates neuronal responses in an asymmetric manner for echolocation and communication context-probe paradigm. At the top, a summary of the experimental results. Direction of the significant changes is indicated with arrows on three different results regarding context-probe paradigm. Note that the changes occur in an asymmetrically between contexts, which can be explained either by **A**, differences in the temporal pattern of the contexts or **B**, differences in the frequency composition of the contexts.

First, we noted that the echolocation and communication sequences used as context differ in their temporal pattern and frequency composition. Therefore, we reasoned two possible scenarios that could explain the observed asymmetries: (i) anaesthesia affects the general neuronal dynamics in a way that only have consequences in the firing rate associated to one context due to its particular temporal modulation (Fig. 7A); or (ii) anaesthesia affects the synaptic inputs associated only to one frequency channel (high-frequency, for example) and therefore it has repercussions in one context due to its carrier frequency (Fig. 7B).

To determine if the asymmetries arose due to differences in the temporal patterns or to the frequency composition of the contexts, we performed a new set of *in vivo* experiments. We built artificial context sequences combining aspects of the original sequences. The new “chimera” sequences included: (i) a “fast echolocation” context, which consisted in several echolocation pulses following the temporal modulation of the original communication context (Fig. 8A left), and (ii) a “slow communication” in which a sequence of distress syllables followed the temporal pattern of the original echolocation context (Fig. 8A right). The probes and the gaps remained unchanged.

**Figure 8.**
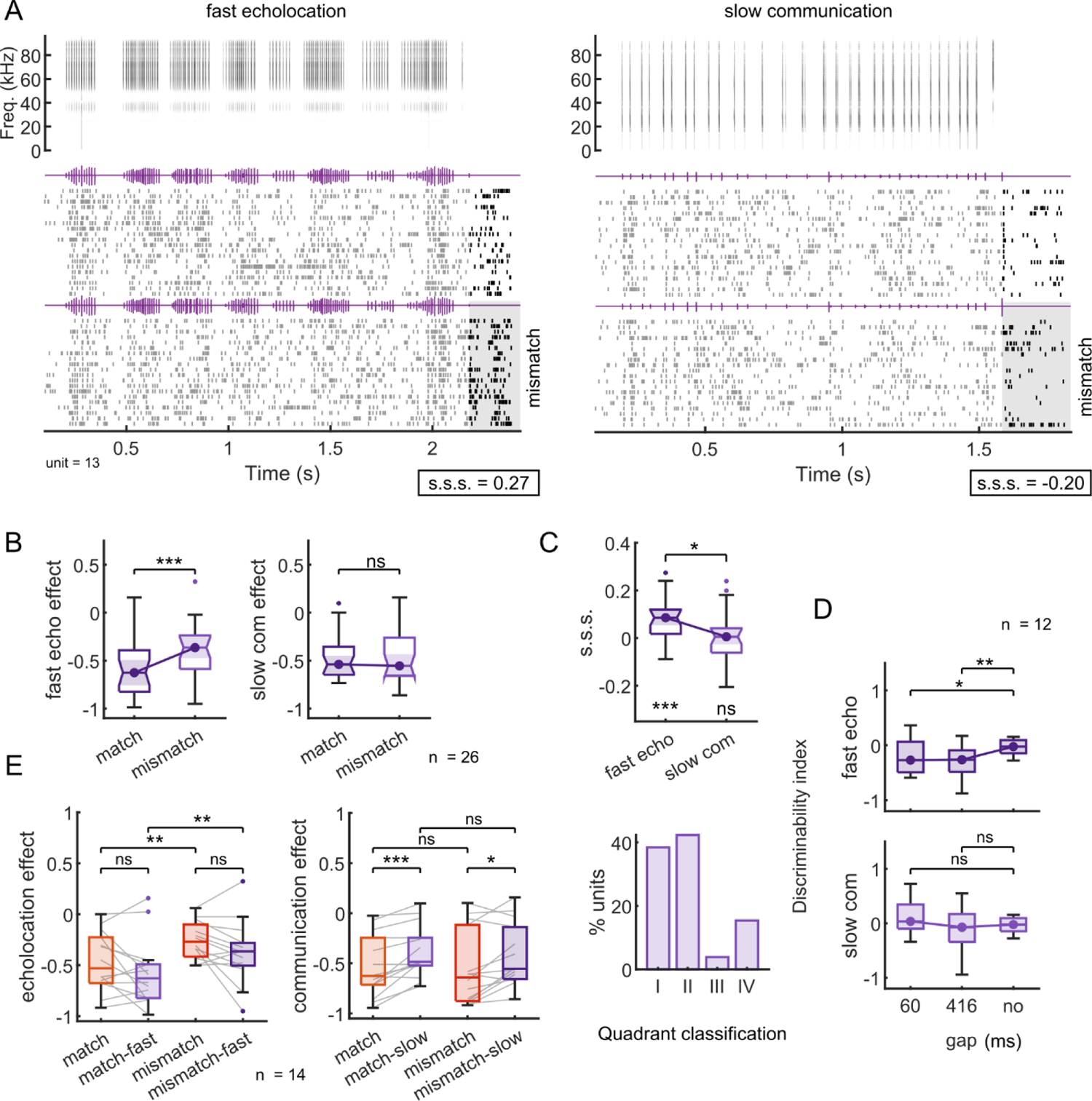
Switching the temporal patterns between contexts does not change the effect on probe-responses under anaesthesia. **A**. Representative example unit firing in response to the context-probe paradigm using “chimera” sequences. Top left, spectrogram of a sequence of echolocation pulse following the temporal pattern of the communication call used as context (called ‘fast echolocation’). Top right, spectrogram of a sequence of communication calls with the temporal pattern of the echolocation sequence used as echolocation context (called ‘slow communication’). Below, the spiking responses to the respective sounds (context) followed by a matching probe sound (top 20 trials) and by a mismatching probe sound (bottom 20 trials). Probe sounds correspond to a single echolocation pulse and a communication call, respectively. The grey square is aligned to the onset of the last call that correspond to the mismatching probe sound in both cases. For each context, the respective value of stimulus-specific suppression (‘s.s.s.’) is indicated. **B**. Context effect after fast echolocation context (left) and after slow communication (n=26). **C**. Top, stimulus-specific suppression (‘s.s.s.’) per context. Significance level below the boxplot indicates differences against null distribution (one sample Wilcoxon signed rank test). Bottom, percentage of units per quadrant as showed in Fig. 3A-B but for units recorded in response to stimuli showed in Fig. 7A. **D**. Discriminability index calculated after 60 and 416 ms from the offset of the fast echolocation context (top) and slow communication (bottom) and after no context. **E**. Same units’ comparison (n=14) between natural temporal pattern context effect (red) and switched temporal pattern context effect (purple). Left, context effect after echolocation context and right, after communication context. All significance levels were obtained using the paired test Wilcoxon signed rank.

We recorded 64 neurons under anaesthesia in response to the artificial context-probe sequences. A total of 26 neurons satisfied the same electrophysiological criteria of being “equally responsive” to both probe sounds in silence. An example is showed in Fig. 8A, the neuron showed preference for the mismatching sound after a sequence of echolocation pulses that doubled the original call rate. However, the neuron did not present a preference for any of the probes after a “slow communication” context.

At the population level, despite the modification of temporal patterns of the calls, the results were similar to those obtained with the natural sequences under anaesthesia. We observed a stronger suppression on the matching probe responses only after “fast echolocation” (Fig. 8B, Wilcoxon signed rank test, *p* = 3.9e-4 for fast echolocation and *p* = 0.9 for slow communication). The “slow communication” context broadly suppressed the responses to both probes, independently on the type of the probe. In agreement with this, the s.s.s. was higher for “fast echolocation” in comparison with “slow communication” (Fig. 8C top, Wilcoxon signed rank test, *p* = 0.03) and the latter exhibited a distribution centred at zero (Fig. 8C top, one sample Wilcoxon signed rank test, *p* = 3.9e-4 for fast-echolocation and *p* = 0.9 for slow communication). The majority of the neurons presented a positive s.s.s. for “fast echolocation” which corresponded to the quadrants I and II (Fig. 8C bottom), based on the neuronal classification in the s.s.s. coordinates plot (Fig. 5A). In terms of neuronal discriminability between the probes, the “fast echolocation” context increased the discriminability index obtained after both gaps, 60 and 416 ms, compared to the index calculated in isolation (Fig. 8D top, Wilcoxon signed rank test, *p* = 0.04 for 60ms-gap and *p* = 0.009 for 416ms-gap). On the contrary, the “slow communication” context did not increase discriminability significantly, neither at gaps of 60 ms nor 416 ms (Fig. 8D bottom, Wilcoxon signed rank test, *p* = 0.15 for 60ms-gap and *p* = 0.6 for 416ms-gap).

To directly test the effect of temporal modulation on the context effects, we performed neuronal recordings in response to both type of contexts, natural and artificial, in the same sessions (n=14, Fig. 8E). Responses in the same neurons showed that increasing the call rate of the echolocation pulses did not modify significantly the context effects on any of the probe-evoked responses (Fig. 8E left, Wilcoxon signed rank test *p* = 0.1 for matching, *p* = 0.08 for mismatching). As observed in the previous data set using intact sequences (Fig. 6A), the suppression driven by echolocation context was stimulus-specific. Interestingly, this was independent on the call rate of the echolocation pulses (Fig. 8E left, Wilcoxon signed rank test *p* = 0.004 for natural, *p* = 0.005 for artificial). On the other hand, a general reduction of the context effect was observed after the “slow communication” context compared to the original (Fig. 8E right, Wilcoxon signed rank test *p* = 9.7e-4 for matching, *p* = 0.02 for mismatching). In this case, decreasing the call rate of the communication context did not lead to stimulus-specific suppression on the probe responses, as well as with the original sequences of communication.

### An updated model predicts anaesthetics effects on auditory cortex

Altogether, our results using chimera sequences suggest that the asymmetrical modulation by KX on the context effects depends on the components of the sequences used as context, more than on their temporal patterns. KX might be then affecting neuronal properties in a frequency-specific manner (as hypothesized in Fig. 7B), which explain why the nonspecific model effects were not able to predict our findings (Figs. 2-3). Accordingly, we modified the awake model in the same manner described in Figs. 2-3, but assuming that anaesthesia affects only one channel input. We ran several models systematically changing channel-specific parameters: either the firing rate of one input or the respective presynaptic adaptation (Table 1, models 1-4). In addition, we examined how an effect on postsynaptic neuronal adaptation modified the frequency-specific effects (Table 1, models 5-8). All of the models presented contradictory results with our data and could therefore not explain our findings.

**Table 1.** Qualitative comparison between multiple models of frequency-specific anaesthesia effect and data obtained *in vivo*. In all models, anaesthesia effects were associated to either high-frequency (HF) tuned inputs or low-frequency (LF) tuned inputs, but never to both. Frequency-specific effects were implemented as a decrease of the cortical input firing rate (‘input’) or an increase of presynaptic adaptation (‘presyn’), either one of them (1-8) or both (9-12). In addition, some of the models can include an increment of postsynaptic adaptation (‘postsyn’), which is not frequency-specific (5-8 and 11-12).

| model | input<br>LF | input<br>HF | presyn<br>LF | presyn<br>HF | postsyn | #<br>effects | contradictory output with *in vivo* data |
| --- | --- | --- | --- | --- | --- | --- | --- |
| 1 | X |  |  |  |  | 1 | decrement of spike rate during communication and no change during echolocation. |
| 2 |  | X |  |  |  | 1 | decrement of echolocation context effect. |
| 3 |  |  | X |  |  | 1 | increment of stimulus-specific suppression after communication context. |
| 4 |  |  |  | X |  | 1 | increment of stimulus-specific suppression after echolocation context. |
| 5 | X |  |  |  | X | 2 | increment of echolocation context effect. |
| 6 |  | X |  |  | X | 2 | decrement of stimulus-specific suppression after echolocation context. |
| 7 |  |  | X |  | X | 2 | increment of echolocation context effect. |
| 8 |  |  |  | X | X | 2 | increment of echolocation context effect on mismatching probe-response. |
| 9 | X |  | X |  |  | 2 | decrement of spike rate during communication and no change during echolocation. |
| 10 |  | X |  | X |  | 2 | decrement of echolocation effect on mismatch probe-responses and negligible communication effect. |
| 11 | X |  | X |  | X | 3 | increment of echolocation effect on both probe-responses |
| 12 |  | X |  | X | X | 3 | none |

Next, we tested combining both frequency-specific effects: input firing rate and presynaptic adaptation together, as showed in Fig. 3. First, we modified the parameters associated to the low-frequency input (Fig. S6). In these models, the echolocation context effects were not affected by anaesthesia (Fig. S6A), which agrees with our electrophysiological data. However, the communication context effect on the mismatching probe responses did not increase (Fig. S6B right) and neurons only decreased spiking activity during communication context (Fig. S6C), both contradicting the experimental results. We reasoned that in order to increase context effect on mismatching probe responses, it is necessary to implement an increment of the postsynaptic adaptation (see Fig. 2C). Therefore, we ran the same simulations (showed in Fig. S6A-C), but with an enhancement of postsynaptic adaptation relative to the awake model (Fig. S6D-F). Indeed, these new simulations showed an increment of communication context effect on mismatching probe-evoked responses (Fig. S6E). However, echolocation also showed an increment on context effects (Fig. S6D). This was due to the fact that postsynaptic adaptation depends on activity of the inputs once they converge in the cortical neurons, and therefore the effect cannot be input-specific and it affects both contexts. In addition, this last model did not change the fact that anaesthesia modulates in a stronger manner the evoked response during communication than during echolocation (Fig. S6F).

Secondly, we modelled the effects of anaesthesia by modifying the input firing rate and the presynaptic adaptation associated to the high-frequency channel (Fig. 9). The simulations showed that:

i. for a certain parameter space, echolocation context on matching probe responses remained unchanged relative to the awake model (Fig. 9A), due to a compensation of the effects (explained in Fig. 3);
ii. echolocation context on mismatching probe responses showed less suppression compared with the awake model, which is not observed in our data (Fig. 9A right);
iii. communication context slightly increased the suppression on both probe responses (Fig. 9B), due to the presence of high-frequency components in communication sounds; and
iv. the effect on the spiking response during echolocation was stronger than communication (Fig. 9C), which agrees with our data.

**Figure 9.**
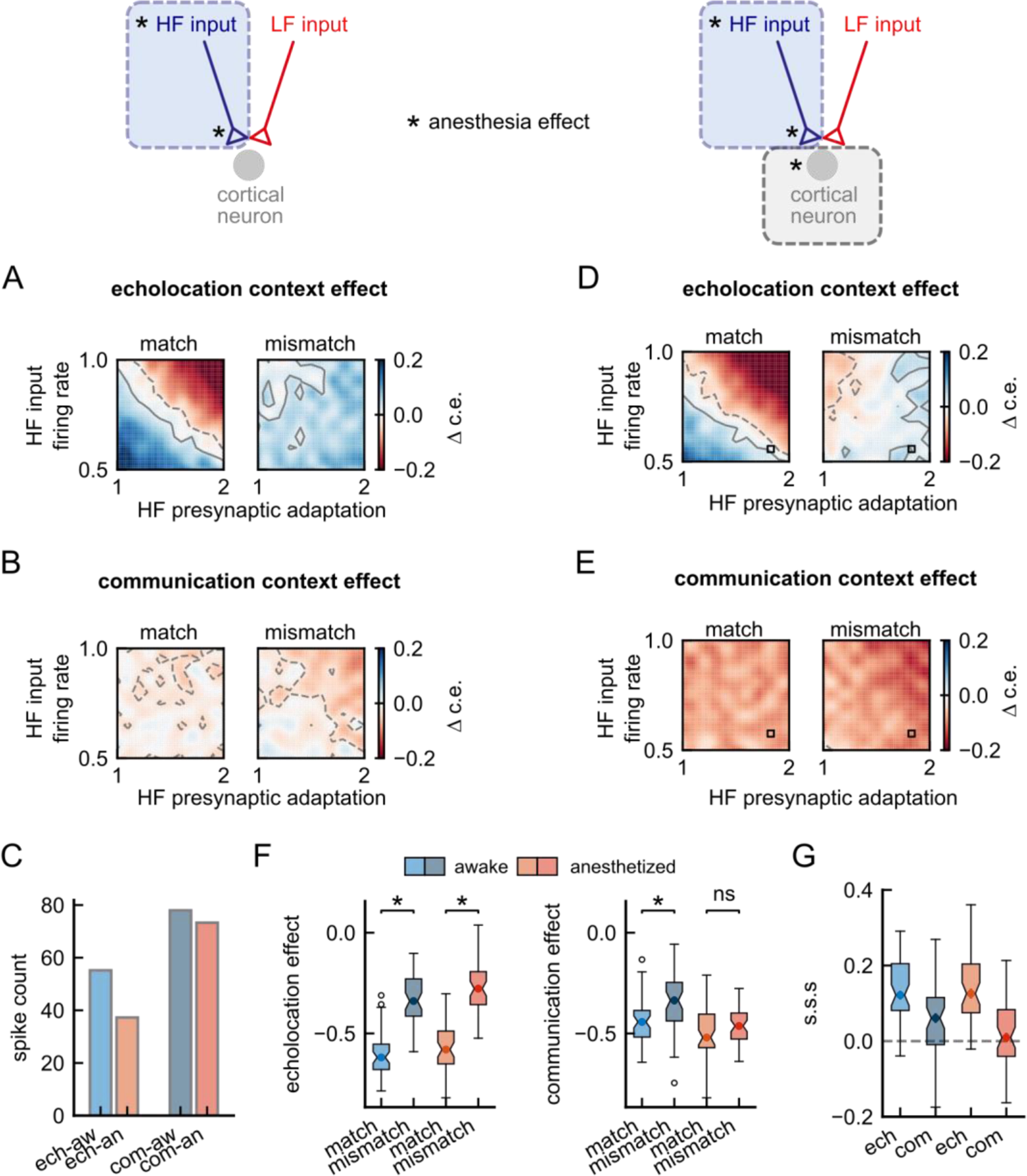
Channel-specific anaesthesia effects on high-frequency cortical inputs can explain the experimental results. **A-C.** Results obtained from an updated model in which anaesthesia affects only high-frequency (HF) cortical inputs, decreasing input firing rate and increasing presynaptic adaptation. **D-G**. Results obtained from a model that includes in addition to those described in ‘A-C’, an increment of postsynaptic adaptation. **A**. Difference between echolocation context effect when decreasing HF inputs firing rate and increasing HF presynaptic adaptation relative to the respective value obtained with the awake model. Contour lines in grey indicate the area with negligible effect size. Positive contours are indicated by solid lines and negative, by dashed lines. The anaesthesia effects were implemented as described in Figure 2. Note that red colours in the colormaps correspond to an increment of suppression, and blue, a decrement. **B**. Same than ‘A’, but difference between communication context effects. **C**. Spike counts during the echolocation context and communication context; for the awake and anesthetized models with LF input firing rate = 0.5 and LF presynaptic adaptation = 1.7. **D-E**. Same than A and B respectively, but for a model that includes postsynaptic adaptation. **F**. Left, context effect after echolocation context, in blue, for the awake model; in red, model under anaesthesia (parameters used are indicated with a square in D and E and correspond to LF input firing rate = 0.5 and LF presynaptic adaptation = 1.7). Right, context effect after communication context. **G**. Stimulus-specific suppression (s.s.s.) per context for model awake and for model anesthetized. The significance levels were obtained using the Wilcoxon rank-sum test.

Interestingly, after combining the described channel-specific effects with an enhancement of postsynaptic adaptation, the outcomes reproduced the totality of our experimental data (Fig. 9D-G). The compensation region of parameters was still present (Fig. 9D left), the echolocation effect on mismatching probe was compensated by the increment of suppression (Fig. 9D right) and communication context suppression on both probes was accentuated (Fig. 9E).

As described above, to reproduce the totality of our experimental data under anaesthesia, we had to change frequency channel-specific parameters in the model in a manner that these parameters compensate each other. To visualize this, we plotted the context effect and s.s.s. for a single combination of parameters within the region of compensation of effects and compared them with the results obtained from the awake model (Fig. 9F-G). The results showed no changes in echolocation context effects between awake and anaesthesia models, with a significant difference between probe responses in both cases (Fig. 9F left, Wilcoxon rank-sum test, *p* = 1.5e-14 for echolocation awake and *p* = 5.7e-15 for echolocation anaesthetized). On the other hand, an increment of suppression was observable on both probe-evoked responses after the communication context with the anaesthesia. As a consequence of this, the significant differences between probe responses disappeared in the anaesthesia model (Fig. 9F right, *p* = 1.0e-4 for communication awake and *p* = 0.09 for communication anaesthetized). Finally, the distribution of s.s.s. was very close to zero only after communication under anaesthesia (Fig. 9G), as observed in our experimental data.

## Discussion

In this study, we recorded cortical activity from anaesthetized bats and compared the responses with those obtained in awake animals. In awake bats, both echolocation and communication acoustic contexts led to stimulus-specific suppression of lagging sounds. However, under anaesthesia, only echolocation context drove stimulus-specific suppression, indicating that anaesthesia has asymmetrical effects on context-dependent processing. We showed that these asymmetries depend on the frequency composition of the sounds used as context, more than on their temporal patterns. We predicted that anaesthesia ‘slows down’ the response dynamics of cortical neurons and has effects on synapses and inputs associated to high-frequency tuned neurons. These effects unbalance cortical inputs and have significant impact on neurons that integrates information across several frequency bands.

### Neuronal mechanisms that underlie anaesthesia effects in the processing of sounds

The neuronal mechanisms affected by anaesthesia during the hearing of natural sounds were studied here by means of computational modelling. The model consisted of a cortical neuron whose inputs were tuned to high or low frequency sounds. Simulations under anaesthesia showed that a reduction in the firing rate of cortical inputs tuned to high-frequency sounds is necessary to reproduce the data obtained *in vivo*.

A frequency-specific decline of neuronal activity could arise from a number of sources, including both the central and the peripheral auditory system. For example, a previous study in the same bat species used here showed that KX reduced inner ear sensitivity preferentially at high frequencies, presumably as a consequence of a metabolic influence on the cochlear amplifier caused by ketamine (Schlenther *et al*., 2014). Similar mechanisms may explain age-related hearing loss in an energy-starved cochlear amplifier (Schmiedt *et al*., 2002). These possible changes in cochlear physiology under anaesthesia may affect central neuron responses, mainly to faint high-frequency sounds, and their transmission to cortical regions.

In agreement with the latter, here we showed that KX significantly reduces the cortical responses to the echolocation context, which consisted of a sequence of several biosonar pulses and their respective faint echoes. Although the decrement of the input activity may be inherited from subcortical areas, the predicted increase of presynaptic adaptation is likely a cortical process. Our model suggested that anaesthesia enhances synaptic depression, predominately in high-frequency tuned inputs. To our knowledge, frequency-specific synaptic effects of anaesthesia have not been reported in the cortex. We speculate that heterogeneous synaptic properties and/or heterogeneous distribution of synapses in these multifunctional neurons may explain a differential gain in the effects observed under anaesthesia.

Anaesthesia also perturbed input-unspecific adaptation in the neuron model, by increasing the time constant of the postsynaptic adaptation process. Consequently, the anaesthetized model had slower dynamics of adaptation than the awake, resulting in a model with slower recovery of the suppressive effects after repeated stimulation during context sequences. Consistent with this, our experimental results showed that the recovery of the context effect was slower under anaesthesia (comparisons between gaps, Fig. 6D-E). Previous studies have shown that, in general, anaesthetics increase the duration of the suppression by preceding stimulus in the auditory cortex (Rennaker *et al*., 2007; Ter-Mikaelian *et al*., 2007; Huetz *et al*., 2009; Scholes *et al*., 2015).

### Anaesthesia unbalances inputs in multifunctional neurons in the auditory cortex

By comparing cortical evoked responses in neurons that respond equally well to echolocation and communication sounds in awake versus anaesthetized bats, we demonstrated that the effects of KX cannot be generalized for both type of vocalizations. In particular, the responses to sounds preceded by communication calls were more affected by anaesthesia than those preceded by echolocation pulses. These asymmetrical effects between echolocation and communication neural processing can be explained by unbalanced synaptic inputs caused by anaesthesia. As their name indicates, multifunctional neurons integrate inputs across multiple frequency bands and their responses to natural sounds result from complex interactions between converging pathways. Other studies have also argued that anaesthesia unbalances synaptic inputs in the cortex (Populin, 2005; Vizuete *et al*., 2012; Zhou *et al*., 2014). Here, we demonstrated that the effects of anaesthesia do not reduce uniformly the neurons responsiveness, but interact in a complex manner to produce non-trivial effects on the spiking responses, which are ultimately measured as readout of cortical activity. Indeed, we showed that a compensation between several effects of KX may underlie seemingly unaffected neuronal responses in the frame of context-dependent processing (Fig. 3). Particularly, we showed this might occur in the processing of vocalizations after echolocation pulses (Fig. 9D). These results highlight the importance of using computational modelling approaches to understand mechanisms that underlie multi-factor processes.

Although our model explained the results by means of excitatory synapses, it is likely that these cortical neurons also receive inhibitory inputs. Ketamine can block glutamatergic receptors associated to inhibitory neurons causing disinhibition (Miller *et al*., 2016). The combination of excitatory and inhibitory inputs and its unbalance under anaesthesia could explain contradictory results and the variability of effects found in the processing of vocalizations across several studies (Capsius & Leppelsack, 1996; George *et al*., 2004; Syka *et al*., 2005; Huetz *et al*., 2009; Schumacher *et al*., 2011; Karino *et al*., 2016). We propose that the variability of anaesthetics effects on evoked responses can be explained by the diversity of synaptic innervation across neurons.

### Cortical processing of natural sounds under anaesthesia

Besides anaesthesia showed asymmetrical effects on the stimulus-evoked activity in our recordings, the effects were solely suppressive. The reduction of spiking was observed during the presentation of long sequences of vocalizations (context sounds) as well as in the context-dependent processing of short calls (probe sounds). Albeit enhancement and no-effect have been reported under anaesthesia in other animals (Capsius & Leppelsack, 1996; Cardin & Schmidt, 2003), suppression seems to be the most frequent outcome in the processing of natural sounds (Vicario & Yohay, 1993; Syka *et al*., 2005). Interestingly, studies using artificial sounds in anaesthetized animals reported stronger activation of neurons in response to sounds than in awake conditions (Zurita *et al*., 1994; Deane *et al*., 2020). The latter could occur due to disinhibition of the cortex, e.g. suppression of inhibitory interneurons. Animal vocalizations are mixtures of time-varying frequencies and amplitudes. Therefore, different patterns of synaptic activation in response to natural sounds and to single pure tones at best frequencies may explain the contradictory effects of anaesthetics.

In our study, we showed that anaesthesia particularly affects context-dependent processing of sounds. The contexts used here were repetitive patterns of shorts sounds, either echolocation pulses or communication syllables, that last a few seconds. A previous study using repetitive broadband sounds showed that responses to clicks in isolation were not affected by anaesthesia, unlike the responses to trains of clicks (Rennaker *et al*., 2007). Also, studies in the same bat species used here demonstrated poor tracking of individual communication syllables under anaesthesia, but good temporal tracking in awake animals (Hechavarria *et al*., 2016b; Martin *et al*., 2017; Garcia-Rosales *et al*., 2018). This suggests that, consistent with our modelling results, the mechanisms underlying temporal processing are affected by ketamine and consequently, the spiking of the neurons after KX injection changes drastically compared to the awake state.

Taken together, the results presented in this article indicate that, while useful, KX anaesthesia has frequency-specific effects in the case of auditory processing of complex sounds, such as vocalizations. Consequently, the effects on stimulus-evoke responses are difficult to predict without combining computational modelling and controlled *in vivo* experiments.

## Methods

### Electrophysiological recordings

Recordings were conducted in 14 bats (10 males) taken from a breeding colony at the Institute for Cell Biology and Neuroscience at the Goethe University in Frankfurt am Main, Germany. Surgery and electrophysiological recordings were performed following the same procedures and experimental setup described elsewhere (Lopez-Jury *et al*., 2021). All experiments were conducted in accordance with the Declaration of Helsinki and local regulations in the state of Hessen (Experimental permit #FU1126, Regierungspräsidium Darmstadt).

To perform the surgery, bats were anaesthetized with a mixture of ketamine (100 mg/ml Ketavet; Pfizer) and xylazine hydrochloride (23.32 mg/ml Rompun; Bayer). Under deep anaesthesia, the dorsal surface of the skull was exposed. The underlying muscles were retracted with an incision along the midline. A custom-made metal rod was glued to the skull using dental cement to fix the head during electrophysiological recordings. After the surgery, the animals recovered for at least 2 days before participating in the experiments.

On the first day of recordings, a craniotomy was performed using a scalpel blade on the left hemisphere in the position corresponding to the auditory region. Particularly, the caudoventral region of the auditory cortex was exposed, spanning primary and secondary auditory cortices (AI and AII, respectively), the dorsoposterior field (DP) and high-frequency (HF) fields. The location of these areas was made following patterns of blood vessels and the position of the pseudocentral sulcus (Esser & Eiermann, 1999; Hagemann *et al*., 2010). Bats were head-fixed and positioned in a custom-made holder over a warming pad whose temperature was set to 27°C. A dose of 0.03 ml of the Ketamine-Xylazine (same concentration used in the surgery) per 20 g of body mass was injected subcutaneously at the beginning of the recording session. In addition, local anaesthesia (Ropivacaine 1%, AstraZeneca GmbH) was administered topically. Each recording session lasted a maximum of 4h. Experiments were made with at least 1-day recovery time in between. In all bats, recordings were performed over a maximum of 14 days.

All experiments were performed in an electrically isolated and sound-proofed chamber. For neural recordings, carbon-tip electrodes (impedance ranged from 0.4 to 1.2 MΩ) were attached to a preamplifier of DAGAN EX4-400 Quad Differential Amplifier system (gain = 50, filter low cut = 0.03 Hz, high cut = 3 kHz). A/D conversion was achieved using a sound card (RME ADI-2 Pro, SR = 192 kHz). Electrodes were driven into the cortex with the aid of a Piezo manipulator (PM 10/1; Science Products GmbH).

Single-unit auditory responses were located at depths of 266 μm +-60 (mean +-SD), using a broadband search stimulus (downward frequency modulated communication sound of the same bat species) that triggered activity in both low and high frequency tuned neurons of layers III-IV in the auditory cortex.

### Acoustic stimulation

We used bat vocalizations to trigger neural activity during the neuronal recordings. The natural sounds were obtained from the same species in previous studies from our lab (Beetz *et al*., 2016; Hechavarria *et al*., 2016a). Acoustic signals were digital-to-analog converted with an RME ADI.2 Pro Sound card and amplified by a custom-made amplifier. Sounds were then produced by a calibrated speaker (NeoCD 1.0 Ribbon Tweeter; Fountek Electronics, China), which was placed 15 cm in front of the bat’s right ear. The speaker’s calibration curve was calculated with a Brüel & Kjaer microphone (model 4135). In addition to the natural sounds, we presented pure tones to the animals to determine the iso-level frequency tuning of each neuron recorded. The pure tones had a duration of 20-ms with 0.5 ms rise/fall time. They were presented randomly in the range of frequencies from 10 to 90 kHz (5 kHz steps, 20 trials) at a fixed sound pressure level of 80 dB SPL. The inter-tone interval was 500 ms.

We studied context-dependent auditory responses using the paradigm described in our previous work (Lopez-Jury *et al*., 2021). Two types of acoustic contexts were presented before probe sounds. The contexts were sequences of echolocation pulses (Fig. S1A) and sequences of distress calls, called “communication” throughout the text (Fig. S1B). The probe sounds were a single echolocation pulse and an individual distress syllable (Fig. S3C). The time interval between the context offset and probe onset was either 60 ms or 416 ms. Therefore, a total of 8 context-stimuli (2 contexts x 2 probes x 2 gaps) were randomly presented and repeated 20 times to the bats during the electrophysiological recordings. In addition, each probe was presented in the absence of context, i.e. after 3.5 s of silence and repeated 20 times as well.

To study how the temporal modulation of context sequences affected our results, we modified the temporal pattern of the natural sequences used as context. These artificial “chimera” sounds corresponded to sequences of echolocation calls following the temporal pattern observed in the natural distress sequence and vice versa. To sweep the temporal pattern of the contexts, we first determined the onset of each call within the sequences as the time at which the envelope of the acoustic signal, measured using the secant method, surpassed a threshold of 70 times the standard deviation above the mean of a silent period of the signal. The silent interval was manually determined, and it corresponded to 260 ms in the communication sequence and 54 ms in the echolocation sequence. The secant method was performed with a MATLAB function called *env_secant* (Martin, 2022) using a window length of 200 for communication and 50 points for echolocation. To avoid false call-onset detections, we set a refractory period of detection of 5 ms and 0.8 ms, respectively. Secondly, each call within the natural sequences was replaced by the calls of the other sequence, keeping the same duration and root-mean-square (RMS) as the original. The RMS was adjusted to match each call-to-call interval. Finally, a calibration was performed with a microphone (model 4135; Brüel & Kjaer) in order to ensure that the chimera sequences had the same RMS than the original natural sounds.

### Paired data sets

In order to directly compare the effects observed between different preparations, we performed separate experiments in which the same neurons were recorded in the two different conditions: (i) in response to natural and constructed contexts and (ii) awake and under anaesthesia. Once we found an auditory neuron, the natural and the constructed contexts were randomly presented after the probes using a gap of 60 ms. We excluded stimuli with longer gaps of 416 ms under this condition since the goal was only to corroborate the previous results, obtained across different sets of neurons. Next, we directly tested the effect of ketamine-xylazine on the neuronal activity. For this purpose, before the recording sessions, we inserted and fixed a cannula subcutaneously in the lower back of the animal. We performed the neuronal recordings in the awake animal and once the recording was finished, we injected the Ketamine-Xylazine mixture (same concentration than the one used in anaesthetized recordings) via the cannula without opening the recording chamber. To monitor the state of the animal, we placed a ceramic piezo vibration sensor (DollaTek) between the bat abdomen and our custom-made holder. This sensor allowed us to record chest movements, as a measure of the respiration rate of the bat. The recordings under anaesthesia were performed only when the respiration rate was steady, lower in amplitude respective to the awake condition and in the absence of animal movements (Fig. S5 top). We were able to record 10 neurons in a total of 5 animals.

### Data analysis

All the recording analyses, including spike sorting, were made using custom-written Matlab scripts (R2018b; MathWorks). The raw signal was filtered between 300 Hz and 3 kHz using a bandpass Butterworth filter (3^rd^ order). To extract spikes from the filtered signal, we detected negative peaks that were at least three standard deviations above the recording baseline; signal in the times spanning 1 ms before the peak and 2 ms after were considered as one spike. The spike waveforms were sorted using an automatic clustering algorithm, “KlustaKwik,” that uses results from PCA analyses to create spike clusters (Harris *et al*., 2000). For each recording, we considered only the spike cluster with the highest number of spikes.

### Electrophysiological classification of units

Neurons were considered as auditory if the number of spikes was above the 95% confidence level calculated for spontaneous firing for the same unit (calculated along 200 ms before the start of each trial) for at least 8ms after the onset of any probe sound in isolation. All the 149 auditory neurons recorded in the current study were classified in terms of the shape of their iso-level frequency tuning curves and in terms of their responses to the probe sounds in isolation (echolocation pulse and distress syllable). The frequency tuning curves were either single-peaked or multipeaked. In order to classify them, the spike count within a time window of 20 ms after the onset of each pure tone was averaged across trials and rescaled by minimum/maximum normalization between the tested frequencies. Single-peaked units were defined as units whose normalized response was higher than or equal to 0.6 only to one half of frequencies measured: either lower than 50kHz (low-frequency tuned) or higher than 50 kHz (high-frequency tuned). On the contrary, multipeaked neurons were those in which the normalized response exceeded 0.6 in both frequency bands (<50 and >=50 kHz). For units that were responsive to both probe sounds in isolation, we categorized them by their preference. Neuronal preferences were determined by comparing the distributions of spikes count during 50 ms after the probes onset, using a non-parametric effect size metric: Cliff’s delta. The present study focused exclusively on those units that presented a negligible or small effect size (abs(Cliff’s delta)<=0.3), called ‘equally responsive’ units. These are units that responded equally well to communication and echolocation sounds presented in isolation. This type of units was previously characterized by us, using the same criterion mentioned above (Lopez-Jury *et al*., 2021). The rest of the neurons presented either preference for one of the probe sounds (abs(Cliff’s delta)>0.3) or unique response to one sound and were excluded from the current study. The number of units per classification; namely ‘equally responsive’, ‘preference to communication’, ‘preference to echolocation’, ‘echolocation only’ and ‘communication only’, is plotted in Fig. S7. The classification was grouped by experimental condition: awake, anaesthetized in response to natural sounds, anaesthetized in response to artificial-natural sounds, anaesthetized in response to both, natural and artificial-natural sounds and before and after anaesthesia injection in response to natural sounds.

### Quantification of context-dependent processing

The effects of the leading acoustic context on probe-triggered responses were quantified by the indices: ‘context effect’, ‘stimulus-specific suppression’ and ‘discriminability index’.

Context effect: quantifies the effect of context (whether echolocation or communication) on the response to a lagging probe sound. This index was calculated as follows: *(R_C,p_ − R_NC,p_)⁄ (R_C,p_ + R_NC,p_),* where *R_C,p_* and *R_NC,p_* corresponds to the number of spikes during 50 ms after the onset of the probe *p* followed by the context *C* and followed by 3.5 s of silence (no context, *NC*), respectively.

Stimulus-specific suppression index: was defined as: *(E_C,m_ − E_C,mm_)* ⁄ 2, where *E_C,m_* is the context effect for context *C* and probe *m*, in which *m matches* with context *C* (i.e. echolocation context and echolocation probe or communication context and communication probe) and *E_C,mm_* for context c and probe *mm*, which *mismatches* with the context *C* (echolocation context and communication probe or communication context and echolocation probe). The index gives values between −1 and 1. Positive values indicate stronger suppressive effect of the context on matching probe responses relative to mismatching probe responses.

Discriminability index: corresponds to a non-parametric effect size measure that quantifies the difference between the responses to both probes across trials. It was calculated for each context using the Cliff’s delta statistic (*d*) between the spike counts during 50 ms after each trial presentation of echolocation (*x_i_*) and communication (*x_j_*) probe sounds, as follows:

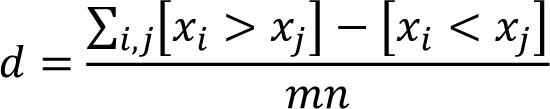

where *m* and *n* are the total number of trials for each probe presentation; and [·] is 1 when the contents are true and 0 when false. According to this, negative Cliff’s delta values indicate higher responses to the communication probe than to the navigation probe. Positive Cliff’s delta values indicate the opposite.

### Computational modelling of anaesthesia effects

We modified an existing model (Lopez-Jury *et al*., 2021) implemented as an integrate-and-fire neuron model. The simulations were run using the Python package Brian2, version 2.3 (Stimberg *et al*., 2019). The model consists of a neuron with subthreshold dynamics that integrates spiking inputs from two narrow frequency channels. The output reproduces the behaviour of the neurons observed in our experiments in awake animals. The synapses are excitatory and exhibit activity-dependent depression. In addition, the neuron presents an adaptive firing threshold that allows it to adapt to inputs that arrives close in time. To study the effects of anaesthesia, we modified three parameters: average input rate ν, synaptic decrement input *Δ_j,s_* and adaptive threshold time constant *τ_th_*. The variation of the parameters was systematically tested by multiplying the original values (awake model) by a vector of equidistant numbers. Therefore, a factor of 1.0 represented the results obtained in the awake model. Throughout the study, the vectors were consistent: for input rate, the vector went from 1.0 to 0.5 in steps of 0.05; in the case of synaptic decrement input, from 1.0 to 2.0 in steps of 0.1 and for varying the adaptive threshold time constant, the vector used went from 1.0 to 2.5 in steps of 0.1.

In the previous model, the inputs to the neuron were modelled by an inhomogeneous Poisson process whose rate is proportional to the envelope of the natural sounds used to stimulate the neurons. The amplitude of the envelope is pondered by the average input rate ν as well as by a factor *k_j,sound_* that represent the responsivity of input *j* to the sound, either echolocation or communication. In order to test the effect of the input rate in the output of the model, we varied the value of *k_j,ech_, k_j,com_* either identically for both inputs (Fig. 2-3) or asymmetrically, only for one input (low-frequency input: Fig. S6 and high-frequency input: Fig. 9). Regarding the increment of presynaptic adaptation by means of the parameter synaptic decrement input *Δ_j,s_*, the variations were either implemented on both inputs (Fig. 3 and Fig. S2) or only for one input (low-frequency input: Fig. S6 and high-frequency input: Fig. 9). Finally, modifications of the adaptive threshold cannot be input-specific since the parameter acts on the postsynaptic neuron. We systematically tested different values using the vector mentioned above (Fig. 2 and Fig. S2) and also, the effect of an increase by a factor of 1.5 on the combination of input rates and presynaptic adaptation effects (Fig. S6 and Fig. 9).

The simulations consisted of 50 neurons, with 20 trials each, in response to the context-probe paradigm. The spiking of the neuron model was analysed exactly as it was done with the electrophysiological data. The spike counts to calculate context effects and responses during contexts were calculated in the same time windows used with the *in vivo* data. Along the study, several anaesthetized models were compared against the original awake model. To quantify the size of the effect on the outputs of the anaesthetized models versus the awake model, we used Cliff’s delta statistics. The effect size was interpreted as negligible, small, medium or large defined by the limits 0.147, 0.333 and 0.474 respectively (Romano *et al*., 2006).

