## Supplementary Figures for "A neuron model with unbalanced synaptic weights explains asymmetric effects of ketamine in auditory cortex"

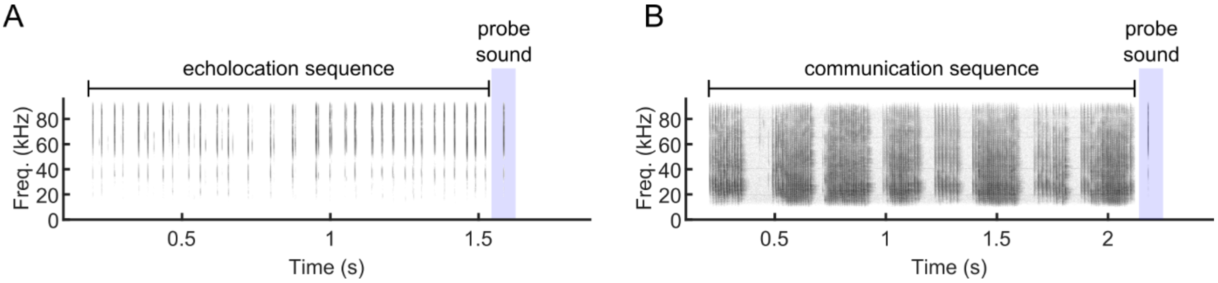


**Figure S1.**

Spectrogram of natural calls of *Carollia perspicillata* associated with echolocation and communication behaviours used to study context dependent sensory processing of natural sounds. **A**. A sequence of echolocation pulses followed by a probe sound, that in this case is a single echolocation pulse. This example corresponds to a 'matching' sound transition. **B**. A distress call followed by a probe sound, i.e. the same echolocation probe used in A. This example corresponds to a 'mismatching' sound transition going from communication (distress) to echolocation.

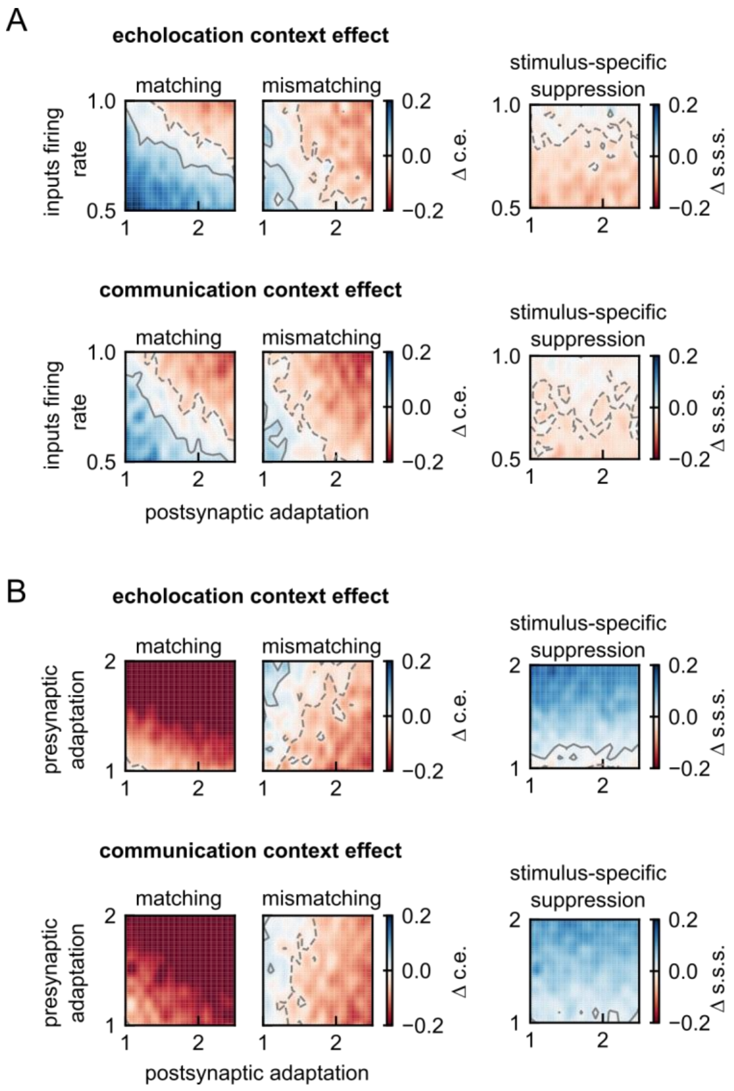


**Figure S2.**

**A**. Difference between context effect (left) and stimulus-specific suppression (right) when decreasing inputs firing rate and increasing postsynaptic adaptation relative to the respective value used in the awake model. First row, after echolocation. Second row, after communication context. The anaesthesia effects were implemented as described in Figure 2. **B**. Difference between context effect (left) and stimulus-specific suppression (right) when increasing presynaptic and postsynaptic adaptation relative to the respective value used in the awake model. First row, after echolocation. Second row, after communication context. The anaesthesia effects were implemented as described in Figure 2. Contour lines in grey indicate the area with negligible effect size. Positive contours are indicated by solid lines and negative, by dashed lines. Note that red colours in the colormaps correspond to an increment of suppression, and blue, a decrement.

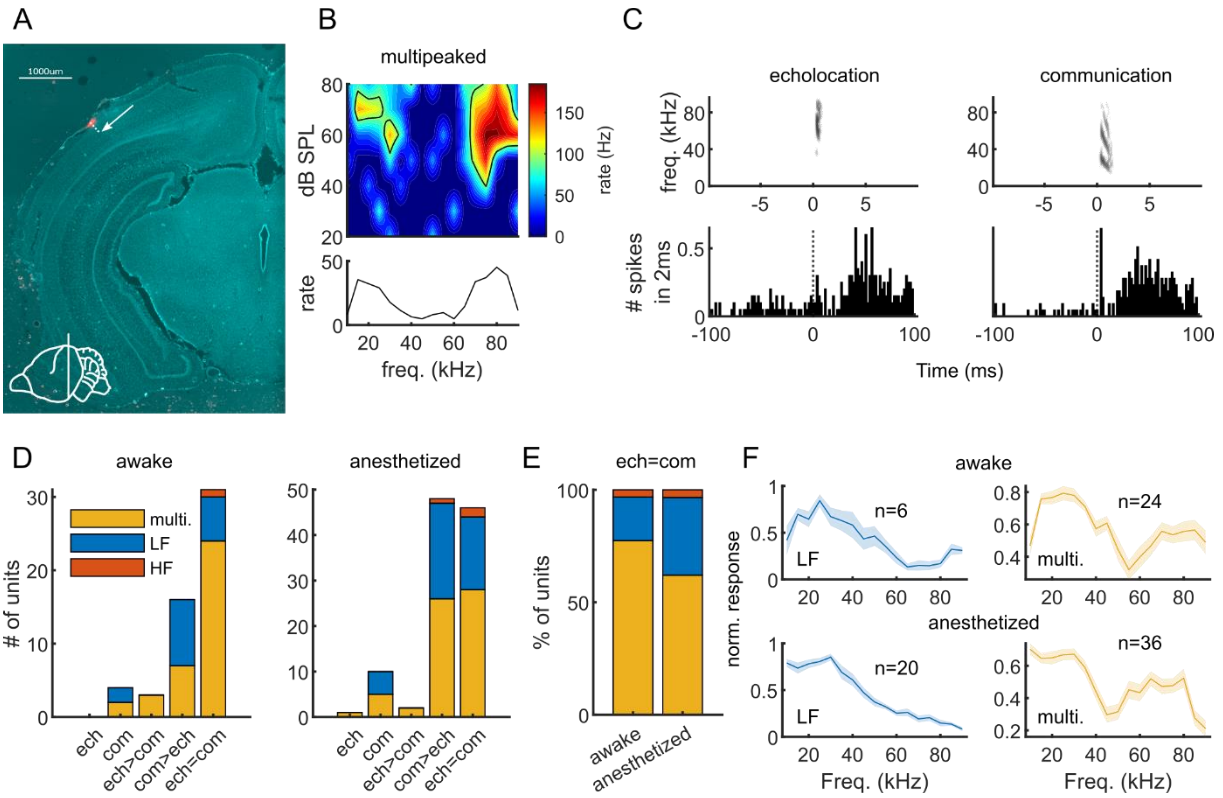


**Figure S3.**

**A**. Coronal section showing the recording site in the bat auditory cortex (AC). The recording enter point was labelled by DiI (red mark) and the recording site calculated with the depth of the electrode (dashed line), indicated by the white arrow. Scale bar indicates 1 mm. The left lower diagram indicates the location of the section in the anterior-posterior axis of the bat brain. **B**. Frequency tuning of a neurons recorded under anaesthesia from the high-frequency fields of bat AC. The firing rate was calculated during 50 ms after the tones' onset. Above, the respective iso-level frequency tuning curve recorded at 80 dB SPL. **C**. Spectrograms of sounds used as probes: an echolocation pulse (left) and a distress syllable (right), below, the respective neuronal response of a representative multipeaked neuron that responded equally well to both sounds categories. **D**. Number of units classified according to responses to echolocation and communication probes recorded in awake (left) and anesthetized bats (right). Categories were defined considering the number of spikes evoked during 50 ms from the sound onset ['ech=com': abs(Cliff's delta) <= 0.3; 'ech<com' or 'com<ech': abs(Cliff's delta) > 0.3]. Colour code indicate the shape of the iso-level frequency tuning curve obtained at 80 dB SPL per natural sound category. Multi.: multipeaked frequency tuning; LF: low-frequency tuned units; HF: high-frequency tuned units. **E**. Percentage of units per preparation: awake and anesthetized that corresponded to each frequency-tuning shape indicated in 'd'. **F**. Average iso-level frequency tuning curve for units that were classified as low-frequency tuned (LF) and as multipeaked (multi.).

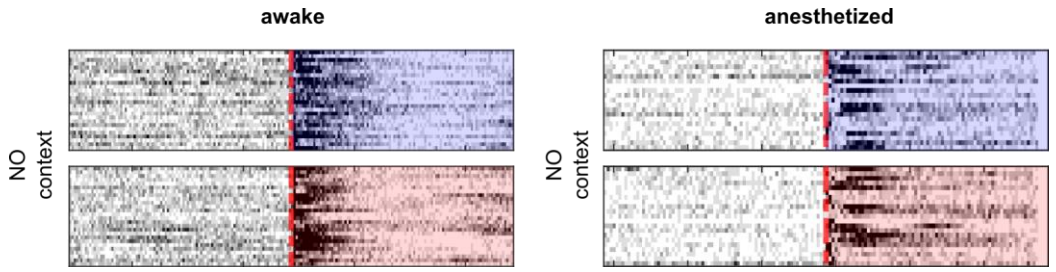


**Figure S4.**

Normalized and averaged firing rate of units that corresponded to the most representative categories in Figure 5B (quadrant I for awake and quadrant II for anesthetized) aligned by probe onset. Blue background corresponds to echolocation probe and red, to communication. Left, for awake preparation, right, for anesthetized preparation. The responses to the probes correspond to those that were preceded by silence.

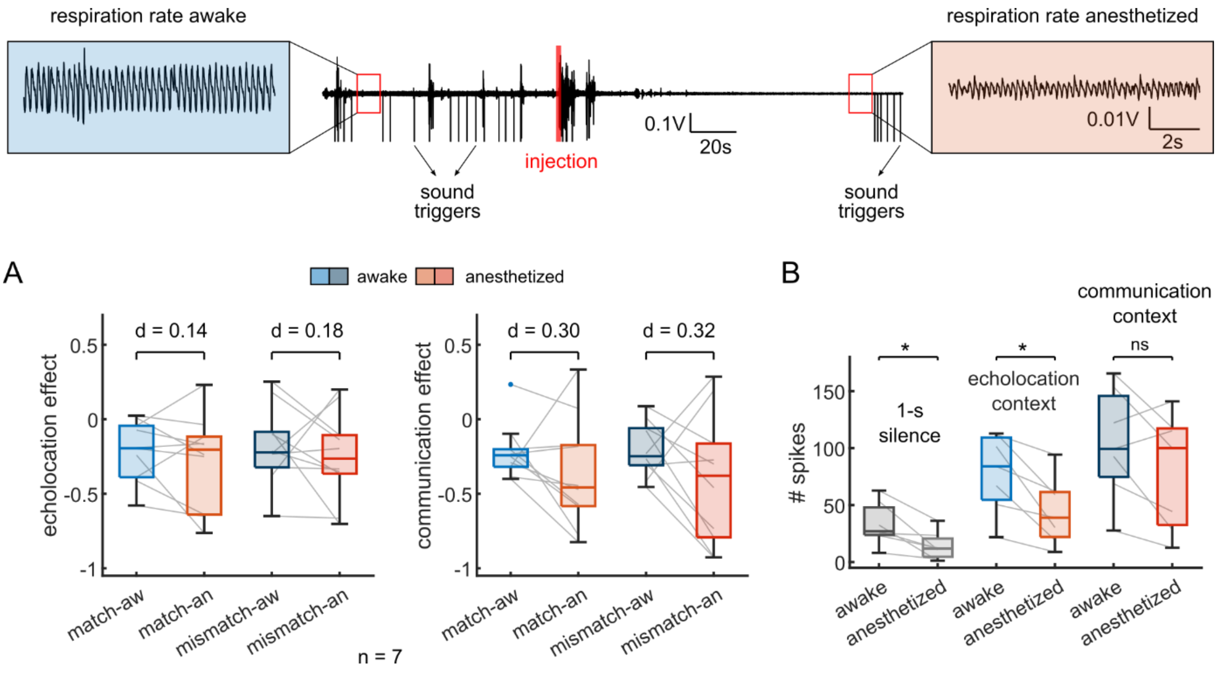


**Figure S5.**

Seven units were recorded during the context-probe paradigm before and after anaesthesia injection. The anaesthesia was injected by using a cannula previously fixed subcutaneously to the animal. The respiration rate of the animal was monitored during the recordings to ensure the success of the anaesthesia injection. Top, black trace shows the piezo signal that monitor the abdominal movements of the bat together with the triggers to natural sounds playbacks. **A**. Context effect on matching and mismatching probe-responses for the same neurons awake and under anaesthesia, after echolocation (left) and communication (right). The comparison before and after were not significant; the cliff's Delta value is indicated on top of the pairs. Grey lines joint the same units. **B**. Spike counts for 1-s of silence, during the echolocation context and communication context; for the same preparation awake and anesthetized. The significance levels were obtained using a paired test (Wilcoxon signed rank).

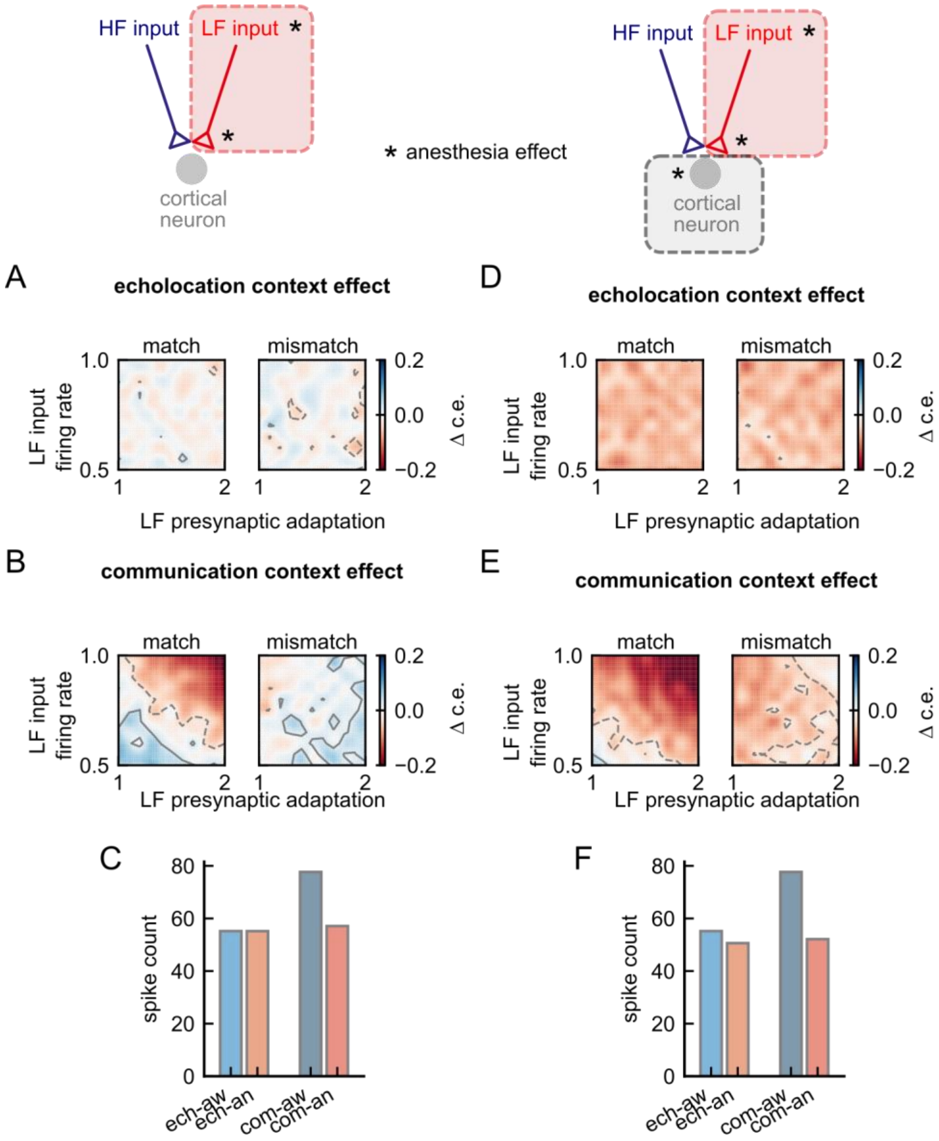


**Figure S6.**

**A-C.** Results obtained from a model in which anaesthesia affects only low-frequency (LF) cortical inputs, as decrement of input firing rate and increment of presynaptic adaptation. **D-F**. Results obtained from a model that includes in addition to those described in ‘A’, an increment in postsynaptic adaptation. **A**. Difference between echolocation context effect when decreasing LF inputs firing rate and increasing LF presynaptic adaptation relative to the respective value obtained with the awake model. Contour lines in grey indicate the area with negligible effect size. Positive contours are indicated by solid lines and negative, by dashed lines. The anaesthesia effects were implemented as described in Figure 2. Note that red colours in the colormaps correspond to an increment of suppression, and blue, a decrement. **B**. Same than ‘A’, but difference between communication context effects. **C**. Spike counts during the echolocation context and communication context; for awake and anesthetized models with LF input firing rate = 0.5 and LF presynaptic adaptation = 1.7. **D-F**. Same than A, B and C respectively, but for a model that include increment of postsynaptic adaptation as an effect of anaesthesia.

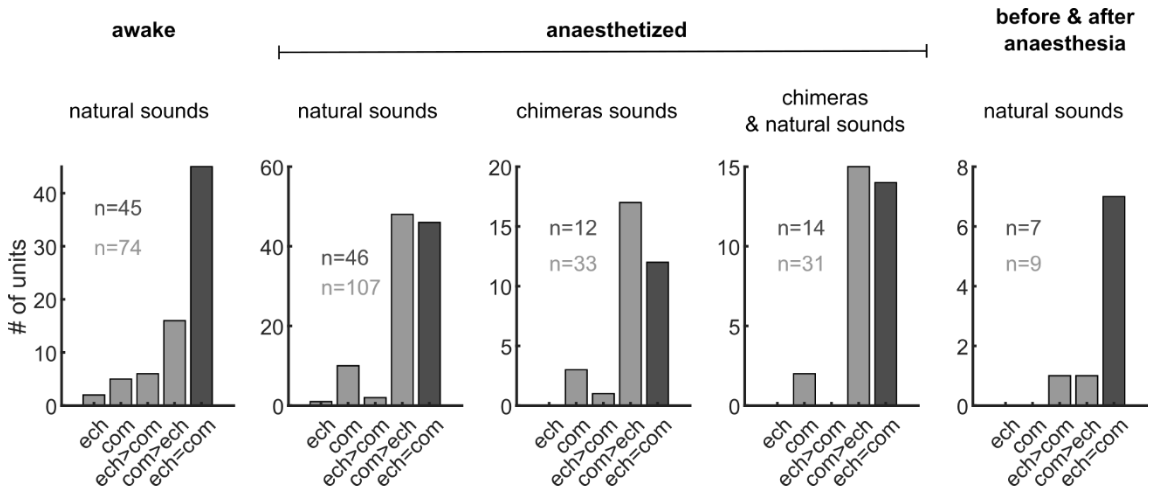


**Figure S7.**

Total number of units recorded for the current study. Note that the analysis presented in this paper only includes those units that were classified as "ech=com", highlighted in black. Natural sounds correspond to the sequences of echolocation and communication showed in Fig S1. Chimeras sounds correspond to sequences of echolocation pulses and distress syllables but after switching the temporal patterns of the original sequences. The last two plots at the right correspond to "paired" data sets. Experiments in which two conditions were tested on the same neurons; either neuronal responses to chimeras sounds versus natural sounds, or neuronal responses in awake and anaesthetized animals.
